# A *Francisella tularensis* L,D-carboxypeptidase plays important roles in cell morphology, envelope integrity, and virulence

**DOI:** 10.1101/817593

**Authors:** Briana Zellner, Dominique Mengin-Lecreulx, Brenden Tully, William T. Gunning, Robert Booth, Jason F. Huntley

**Affiliations:** Department of Medical Microbiology and Immunology, University of Toledo, Toledo, OH, U.S.A.; Université Paris-Saclay, CEA, CNRS, Institute for Integrative Biology of the Cell (I2BC), 91198, Gif-sur-Yvette, France; Department of Pathology, University of Toledo, Toledo, OH, U.S.A.

**Author notes:** To whom correspondence should be addressed: Jason F. Huntley, Department of Medical Microbiology and Immunology, University of Toledo College of Medicine and Life Sciences, 3000 Arlington Ave., MS1021, HEB 200D, Toledo, OH, U.S.A., 43614.

**Keywords:** tularemia, peptidoglycan, L,D-carboxypeptidase, virulence, *Francisella* tularensis

## Abstract

*Francisella tularensis* is a Gram-negative, intracellular bacterium that causes the zoonotic disease tularemia. Intracellular pathogens, including *F. tularensis*, have evolved mechanisms to survive in the harsh environment of macrophages and neutrophils, where they are exposed to cell envelope-damaging molecules. The bacterial cell wall, primarily composed of peptidoglycan (PG), maintains cell morphology, structure, and membrane integrity. Intracellular Gram-negative bacteria protect themselves from macrophage and neutrophil killing by recycling and repairing damaged PG – a process that involves over 50 different PG synthesis and recycling enzymes. Here, we identified a PG recycling enzyme, L,D-carboxypeptidase A (LdcA), of *F. tularensis* that is responsible for converting PG tetrapeptide stems to tripeptide stems. Unlike *E. coli* LdcA and most other orthologs, *F. tularensis* LdcA does not localize to the cytoplasm and also exhibits L,D-endopeptidase activity, converting PG pentapeptide stems to tripeptide stems. Loss of *F. tularensis* LdcA led to altered cell morphology and membrane integrity, as well as attenuation in a mouse pulmonary infection model and in primary and immortalized macrophages. Finally, an *F. tularensis ldcA* mutant protected mice against virulent Type A *F. tularensis* SchuS4 pulmonary challenge.

## Introduction

The Gram-negative bacterial cell wall plays an important role in maintaining cell shape, protecting against external insults, and preventing cell lysis amid fluctuations in internal turgor pressure (Dhar, 2018). The cell wall is composed of peptidoglycan (PG), a network of alternating *N*-acetylglucosamine (GlcNAc) and *N*-acetylmuramic acid (MurNAc) glycan chains that are crosslinked through peptide stems, and lies just outside of the cytoplasmic membrane of most bacteria (Johnson, 2013, Mengin-Lecreulx & Lemaitre, 2005). In *Escherichia coli*, PG has been shown to be covalently attached to the outer membrane (OM) by Braun’s lipoprotein and noncovalently attached by Pal and other lipoproteins (Braun, 1969, Braun, 1975, Bouveret, 1999, Leduc, 1992). A loss of membrane integrity can occur if interactions between PG and attached lipoproteins are disturbed, thus maintenance of correct PG architecture is extremely important (Braun & Rehn, 1969, Braun & Hantke, 2019).

Synthesis of Gram-negative PG precursors, studied mainly in *E. coli*, occurs in the bacterial cytoplasm and requires a series of enzymes to build a pentapeptide PG monomer before transporting this molecule into the periplasm. Periplasmic PG cross-linking most often occurs between the fourth residue (D-Ala) of newly-formed pentapeptide stems to the third residue (*meso*-A_2_pm) of existing tripeptide stems (4-3 cross links), resulting in the release of the pentapeptide terminal D-Ala and forming tetrapeptide stems (Glauner, 1988, Pazos & Peters, 2019). PG is not a static structure, rather, PG degradation/remodeling is necessary to incorporate new PG and expand the cell wall during bacterial growth and replication, to insert flagella or secretion systems, and to septate during bacterial division (Scheurwater, 2008). Up to 60% of PG is recycled per generation, helping to repair damaged PG and providing energy during periods of stress or starvation (Dhar, 2018, Park, 2008, Holtje, 1998). Using *E. coli* as a model, more than 50 different PG synthesis and hydrolysis/recycling enzymes have been identified, most of which appear to be cytoplasmic. While deletion studies in *E. coli* have indicated that some PG synthesis enzymes are essential, few PG hydrolases have been found to be essential and, in fact, substantial redundancy in hydrolase activity appears to exist. Indeed, alterations in cell morphology, membrane integrity, and ability to replicate/divide sometimes are observed only after multiple PG metabolism genes have been deleted (Dhar, 2018, van Heijenoort, 2011, Vollmer & Bertsche, 2008). Given the increasing threat of antimicrobial-resistant organisms and link between PG homeostasis and bacterial virulence, more studies are needed to understand how a diverse range of Gram-negative bacteria synthesize and recycle PG (Juan, 2018). Finally, PG studies can reveal new strategies to treat bacterial infections, as an *Acinetobacter baumannii* penicillin-binding protein (PBP) mutant was reported to be more sensitive to complement-mediated killing than wild-type bacteria (Russo, 2009) and a *Helicobacter pylori* PG hydrolase (AmiA) mutant was unable to colonize mouse stomachs (Chaput, 2016).

PG recycling, mainly characterized in *E*. *coli,* begins with periplasmic lytic transglycosylase (LT) cleavage of the β-1,4-glycosidic linkage between MurNAc and GlcNAc, forming a GlcNAc-1,6-anhydro-MurNAc product, which creates sites for the insertion of PG precursors and recycling of the GlcNAc-1,6-anhydro-MurNAc peptide (Scheurwater, 2008). Low molecular mass penicillin-binding proteins (LMM PBPs) can function as endopeptidases, cleaving the cross-links between adjacent tetrapeptide stems, and/or as D,D-carboxypeptidases, removing the terminal D-Ala of pentapeptides, forming the tetrapeptide (Dhar, 2018). Inner membrane permeases, such as AmpG, then transfer GlcNAc-1,6-anhydro-MurNAc disaccharides, with or without attached peptides, to the cytoplasm where they can be disassembled. Cytoplasmic NagZ and AmpD further degrade disaccharides by cleaving the bond between GlcNAc and 1,6-anhydro-MurNAc, and separating the peptide chain from 1,6-anhydro-MurNAc, respectively. Additional cytoplasmic hydrolases, such as L,D-carboxypeptidases (Ldc), act on free peptide chains to cleave the terminal D-Ala from tetrapeptides, resulting in tripeptides (Dhar, 2018). PG is unusual in that it is both highly dynamic (*e.g.,* allowing for bacterial division and molecular transport across the periplasm), yet tightly regulated to prevent membrane collapse and bacterial death. As such, PG recycling enzymes have been speculated to be important virulence determinants in Gram-negative bacteria (Juan, 2018). Indeed, *E. coli ldc* mutants lyse in stationary phase (Templin, 1999) and are more susceptible to β-lactam antibiotics (Ursinus, 1992), *Helicobacter* and *Campylobacter ldc* mutants have altered cell morphology and defects in motility and biofilm formation (Sycuro, 2013, Frirdich, 2012, Frirdich, 2014), and *Neisseria gonorrhoeae ldc* mutants are unable to stimulate NOD1-dependent responses in the host (Lenz, 2017). However, very little is known about the importance of PG recycling enzymes in the pathogenesis of intracellular pathogens such as *Francisella tularensis*.

*F. tularensis*, the causative agent of tularemia, is a Gram-negative, intracellular, coccobacillus that can infect and cause lethal disease in many species, including humans (Dennis, 2001, Keim, 2007). There are three subspecies of *F. tularensis*, subsp. *tularensis* (Type A), subsp. *holarctica* (Type B), and subsp. *mediasiatica*, although only subsp. *tularensis* and subsp. *holarctica* are virulent for humans (Kingry, 2014). *F. tularensis* poses a severe threat to public health and has been classified as an NIH Category A Priority Pathogen and a CDC Tier 1 Select Agent due to its low infectious dose (<10 CFU), ease of aerosolization, and high morbidity and mortality rates (up to 60%) (Ellis, 2002, Sjostedt, 2007). Like other intracellular pathogens, *F. tularensis* has evolved different mechanisms to infect, survive, and replicate within host cells, including macrophages and neutrophils (Ray, 2009). However, this lifestyle exposes the bacteria to reactive oxygen species (ROS), reactive nitrogen species (RNS), antimicrobial peptides, and other cell membrane- and cell wall-damaging molecules (Jones, 2012). Our group previously demonstrated that the *F. tularensis* disulfide bond formation protein A (DsbA) ortholog repairs damaged outer membrane proteins and known virulence factors. We additionally showed that *F. tularensis* DsbA, unlike periplasmic DsbA in *E. coli* and most other Gram-negative bacteria, is outer membrane-bound and is a multifunctional protein with both oxidoreductase and isomerase activities. Finally, using a molecular trapping approach, we identified over 50 *F. tularensis* DsbA substrates, many of which we speculate are involved in virulence (Ren, 2014).

Here, we determined the function of one of those *F. tularensis* DsbA substrates – a previously unstudied hypothetical protein containing a putative LdcA domain – and assessed its role in bacterial virulence. Deletion of *F. tularensis ldcA* resulted in bacteria with altered cell morphology, increased sensitivity to β-lactam antibiotics, yet increased resistance to several stressors (*e.g.,* H_2_O_2,_ NaCl, low pH). Next, we demonstrated that *F. tularensis* LdcA exhibits L,D-carboxypeptidase and L,D-endopeptidase activities on pentapeptide and tetrapeptide residues of PG. Finally, we established that *F. tularensis* LdcA is required for virulence, as mutants were unable to replicate in macrophages or cause disease in mice.

## Results

### FTL1678 contains a putative L,D-carboxypeptidase domain

Previous studies by our group and others have shown that *F. tularensis* DsbA mutants are attenuated in mice (Qin, 2009, Ren, 2014). However, additional work by our group, demonstrating that DsbA possesses both oxidoreductase and isomerase activities to repair damaged envelope and cell membrane proteins, indicated that other envelope proteins likely are responsible for *F. tularensis* virulence (Ren, 2014). To identify new *F. tularensis* virulence factors, we used a molecular trapping approach and identified over 50 *F. tularensis* DsbA substrates (Ren, 2014). One of those DsbA substrates, FTL1678, is annotated in the *F. tularensis* genome as a conserved membrane hypothetical protein. Here, a conserved domain search revealed that a large portion of FTL1678 contains a putative Ldc domain, part of the peptidase_S66 superfamily (Figure S1). Ldc proteins have been studied in a number of Gram-negative bacteria, including *E. coli* (Metz, 1986a, Templin, 1999, Metz, 1986b, Ursinus, 1992), *Pseudomonas aeruginosa* (Korza, 2005), *N. gonorrhoeae* (Lenz, 2017), and *Campylobacter jejuni* (Frirdich, 2014). To further explore this conserved domain, amino acid sequences of FTL1678 (*F. tularensis* subsp. *holarctica* [Type B] LVS) and FTT0101 (homolog of FTL1678 in *F. tularensis* subsp. *tularensis* [Type A] SchuS4) were aligned with LdcA orthologs from *E. coli*, *aeruginosa*, *N. gonorrhoeae*, and *C. jejuni* (named Pgp2). Despite low percentages of amino acid identities among the LdcA orthologs (6.3% to 30.3%; Figure 1), there was a higher degree of amino acid similarity among LdcA orthologs (13.0% [*E. coli* and *C. jejuni*] to 44.7% [*E. coli* and *N. gonorrhoeae*]; Figure 1). Notably, the LdcA Ser-Glu-His catalytic triad, previously shown to be required for *P. aeruginosa* LdcA activity (Korza, 2005), was absent from *C. jejuni* Pgp2 but was present in all LdcA homologs, including FTL1678 and FTT0101 (Figure 1).

**Figure 1.**
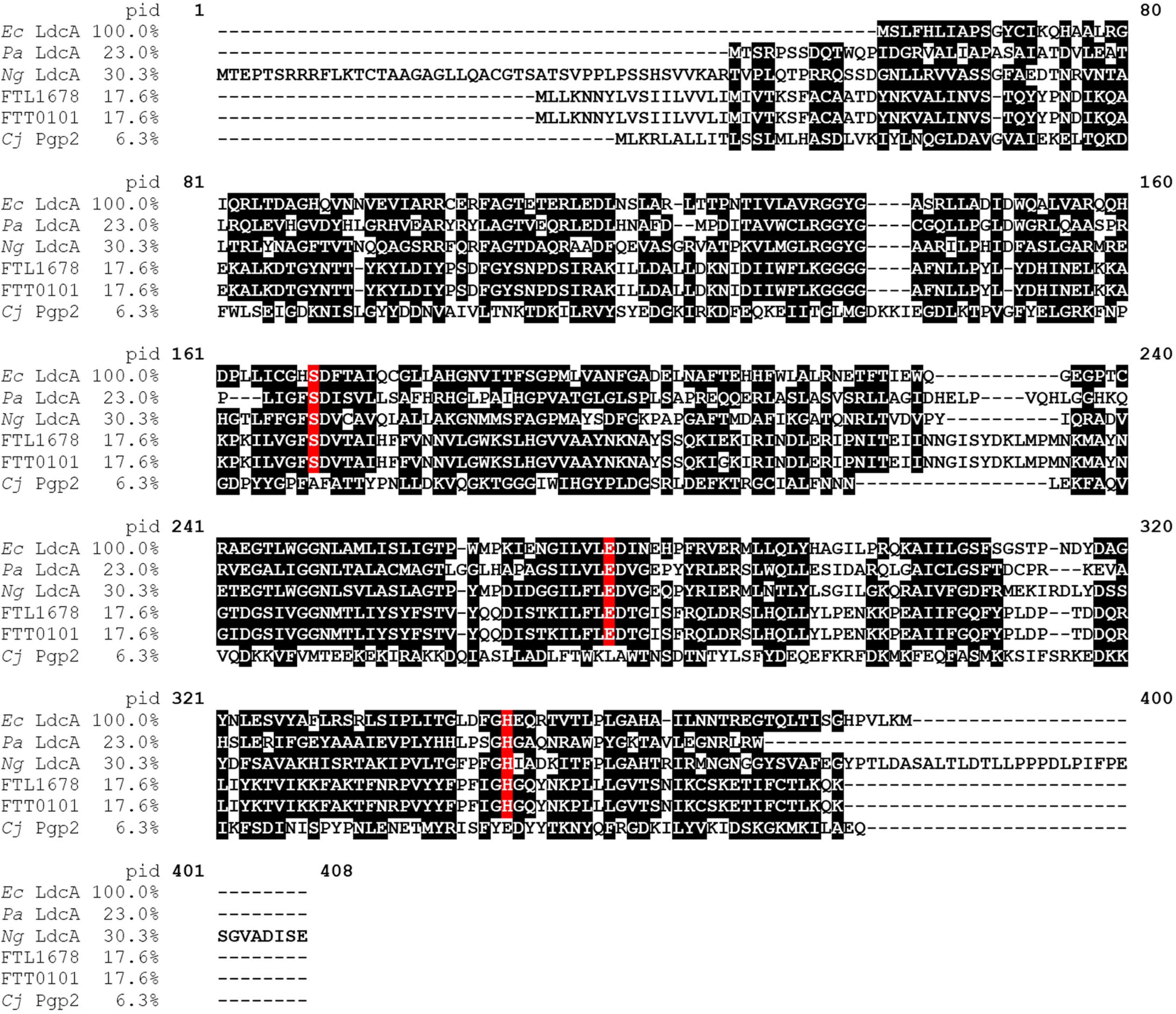
Amino acid alignment of bacterial L,D-carboxypeptidases. Clustal Omega amino acid alignment of *E. coli* LdcA (BAA36050.1), *P. aeruginosa* LdcA (Q9HTZ1), *N. gonorrhoeae* LdcA (YP_208343.1), *F. tularensis* Type B FTL1678, and *F. tularensis* Type A FTT0101, and *C. jejuni* Pgp2 (WP_002856863). Percent identities (pid), compared to *E. coli* LdcA, are indicated. Black shading indicates similar residues. Red shading indicates the catalytic triad.

### FTL1678 exhibits L,D-carboxypeptidase and L,D-endopeptidase activities

To confirm the predicted Ldc activity of FTL1678 and FTT0101, recombinant FTL1678 and FTT0101 were expressed and affinity purified from *E. coli*. As a control, lysates from *E. coli* containing the empty expression vector (pPROEX HTb) also were affinity-purified. PG precursors and PG intermediates were prepared as previously described (Herve, 2007, Blanot, 1983, Pennartz, 2009, Leulier, 2003, Stenbak, 2004). HPLC retention times for all PG substrates and products are listed in Table S1. Recombinant FTL1678, lysate from the vector control, or buffer alone were incubated with various PG substrates (Table 1) to determine substrate specificity and specific activity. The vector control and buffer alone did not demonstrate activity against any of the PG substrates (Figure S2 and S3). When FTL1678 was incubated with various PG substrates, the highest specific activity was detected against the tetrapeptide substrates GlcNAc-anhydroMurNAc-L-Ala-γ-D-Glu-*meso*-A_2_pm-D-Ala (tracheal cytotoxin; TCT; 21.5 nmol/min/mg of protein; Table 1) and GlcNAc-MurNAc-L-Ala-γ-D-Glu-*meso*-A_2_pm-D-Ala (reducing PG monomer; 15.6 nmol/min/mg of protein; Table 1), confirming that FTL1678 exhibits L,D-carboxypeptidase activity. Interestingly, FTL1678 activity against free tetrapeptide, L-Ala-γ-D-Glu-*meso*-A_2_pm-D-Ala, was approx. 6-fold lower (3.4 nmol/min/mg of protein; Table 1) than TCT and 5-fold lower than the reducing PG monomer (Table 1), indicating that GlcNAc and MurNAc may be important for tetrapeptide recognition or FTL1678 binding. Next, FTL1678 was found to exhibit negligible activity against L-lysine-containing substrates (0.7 to 1.3 nmol/min/mg of protein; Table 1), where L-lysine replaced *meso*-A_2_pm at the third amino acid position, indicating the importance of *meso*-A_2_pm. Importantly, FTL1678 exhibited specific activity against pentapeptide substrates MurNAc-L-Ala-γ-D-Glu-*meso*-A_2_pm-D-Ala-D-Ala (9.8 nmol/min/mg of protein; Table 1) and UDP-MurNAc-L-Ala-γ-D-Glu-*meso*-A_2_pm-D-Ala-D-Ala (5.9 nmol/min/mg of protein; Table 1), indicating that FTL1678 also functions as an L,D-endopeptidase (cleavage of the pentapeptide between *meso*-A_2_pm and D-Ala). D-Ala-D-Ala, but not D-Ala, was released in that case, confirming that FTL1678 did not display D,D-carboxypeptidase activity. FTL1678 was not inhibited by 5 mM EDTA and did not require the presence of cations (Mg^2+^) for tetrapeptide cleavage (Figure S3).

**Table 1.** Specific activity and substrate specificity of FTL1678 and FTT0101 enzymes.

| Substrate | Specific activity (nmol/min/mg of protein)<sup>a</sup> |  |
| --- | --- | --- |
|  | FTL1678 | FTT0101 |
| **A<sub>2</sub>pm-containing substrates** |  |  |
| GlcNAc-anhydroMurNAc-L-Ala-γ-D-Glu-*meso*-A<sub>2</sub>pm-D-Ala (TCT) | 21.5 ± 1.34 | 29.0 ± 1.42 |
| GlcNAc-MurNAc-L-Ala-γ-D-Glu-*meso*-A<sub>2</sub>pm-D-Ala (PG monomer) | 15.6 ± 0.79 | 24.2 ± 1.80 |
| TCT dimer (4-3 D,D-crosslinkage) | 7.7 ± 0.30 | 11.8 ± 0.53 |
| MurNAc-L-Ala-γ-D-Glu-*meso*-A<sub>2</sub>pm-D-Ala | 5.8 ± 0.66 | 7.8 ± 1.10 |
| UDP-MurNAc-L-Ala-γ-D-Glu-*meso*-A<sub>2</sub>pm-D-Ala | 4.6 ± 0.52 | 6.3 ± 0.26 |
| L-Ala-γ-D-Glu-*meso*-A<sub>2</sub>pm-D-Ala (free tetrapeptide) | 3.4 ± 0.59 | 8.0 ± 0.92 |
| MurNAc-L-Ala-γ-D-Glu-*meso*-A<sub>2</sub>pm-D-Ala-D-Ala (pentapeptide) | 9.8 ± 1.22 | 6.2 ± 0.56 |
| UDP-MurNAc-L-Ala-γ-D-Glu-*meso*-A<sub>2</sub>pm-D-Ala-D-Ala (pentapeptide) | 5.9 ± 0.14 | 6.4 ± 0.17 |
| **L-Lysine-containing substrates** |  |  |
| MurNAc-L-Ala-γ-D-Glu-L-Lys-D-Ala | 1.3 ± 0.28 | 2.1 ± 0.36 |
| L-Ala-γ-D-Glu-L-Lys-D-Ala | 0.7 ± 0.14 | 2.1 ± 0.28 |
| UDP-MurNAc-L-Ala-γ-D-Glu-L-Lys-D-Ala-D-Ala | 1.2 ± 0.26 | 1.7 ± 0.20 |
| **Others** |  |  |
| GlcNAc-MurNAc-L-Ala-γ-D-Glu-*meso*-A<sub>2</sub>pm(NH<sub>2</sub>)-D-Ala |  | Very low (<0.5) |
| GlcNAc-MurNAc-L-Ala-γ-D-Glu(NH<sub>2</sub>)-*meso*-A<sub>2</sub>pm-D-Ala |  | Not detected<sup>b</sup> |
| GlcNAc-MurNAc-L-Ala-γ-D-Glu(NH<sub>2</sub>)-*meso*-A<sub>2</sub>pm(NH<sub>2</sub>)-D-Ala |  | Not detected<sup>b</sup> |
| Peptidoglycan polymer |  | Not detected<sup>b</sup> |

<sup>a</sup> Standard enzyme assay conditions are described in the Materials and Methods. Values represent the mean ± standard deviation of triplicate experiments.
<sup>b</sup> Not detected indicates that no formation of the corresponding disaccharide-tripeptide (HPLC) and/or release of alanine (amino acid analyzer) was detected in the assay conditions used.

To further investigate the endopeptidase activity of FTL1678, FTL1678 was incubated with various TCT monomer and dimer substrates, containing different peptide lengths and cross-link locations (Table S2). Although FTL1678 exhibited the highest specific activity against TCT monomers containing tetrapeptide stems with either an alanine or a glycine in the fourth position (20.1 nmol/min/mg and 18.0 nmol/min/mg of protein, respectively; Table S2), FTL1678 also was able to cleave all four variations of the TCT dimer (two cross-linked TCT monomers; Table S2). Together, these results demonstrated that FTL1678 exhibited both L,D-carboxypeptidase and L,D-endopeptidase activities. Of the four different TCT dimer analogs tested, FTL1678 was most active on TCT dimers connected by a 4-3 (D-D) D-Ala-*meso*-A_2_pm cross linkage (6.6 nmol/min/mg of protein; Table S2), followed by an approx. 24-fold reduction in activity on 3-3 (L-D) A_2_pm-A_2_pm cross links connecting two tripeptides (0.28 nmol/min/mg of protein; Table S2), a tripeptide and tetrapeptide with a glycine at the fourth position (0.21 nmol/min/mg of protein; Table S2), and a tripeptide and tetrapeptide with an alanine at the fourth position (0.16 nmol/min/mg of protein; Table S2). These data suggest that, while FTL1678 exhibits endopeptidase activity on 4-3 and 3-3 cross-linked dimers, cleavage of 3-3 cross links is likely not its main physiological function.

**Table 2.** Sensitivity of WT and Δ*FTL1678* to antibiotics, detergents, and dyes.

| Compound | Concentration ($\mu\text{g}/\text{disk}$) | Average zone of inhibition, mm (mean ± SD)<sup>a</sup> | |
| --- | --- | --- | --- |
| | | WT | $\Delta FTL1678$ |
| Gentamicin | 4 | 3 ± 0 | 1 ± 0 (R) |
| Tetracycline | 5 | 2 ± 0 | 1 ± 0 (R) |
| Chloramphenicol | 5 | 3 ± 0 | 1 ± 0 (R) |
| Ciprofloxacin | 5 | 5 ± 0 | 2 ± 0 (R) |
| Ampicillin | 200 | 3 ± 0 | 4 ± 0 (S) |
| Vancomycin | 20 | 1 ± 0 | 3 ± 0 (S) |
| Bacitracin | 182 | 1 ± 0 | 1 ± 0 |
| Polymyxin B | 100 | 1 ± 0 | 1 ± 0 |
| Lysozyme | 1000 | 1 ± 0 | 2 ± 0 (S) |
| Ethidium Bromide | 5 | 3 ± 0 | 1 ± 0 (R) |
| Triton-X100 | 750 | 3 ± 0 | 3 ± 0 |
| SDS | 1000 | 1 ± 0 | 2 ± 0 (S) |
| CTAB | 50 | 1 ± 0 | 1 ± 0 |
| CHAPS | 50 | 1 ± 0 | 1 ± 0 |

<sup>a</sup> (R) indicates $\Delta FTL1678$ is significantly more resistant than WT by one-way ANOVA ( $P < 0.05$ )
<sup>a</sup> (S) indicates $\Delta FTL1678$ is significantly more sensitive than WT by one-way ANOVA ( $P < 0.05$ )

Assays were repeated with recombinant FTT0101 (SchuS4 homolog) and, due to 99.4% amino acid identify with FTL1678, FTT0101 demonstrated similar tetrapeptide cleavage activity (*i.e.,* LdcA activity) as FTL1678 (Table 1). FTT0101 also was not active on a peptidoglycan polymer and had either no or negligible activity on PG monomers that were amidated at the *meso*-A_2_pm or D-Glu residues (Table 1).

As highlighted in Figure 1, a Ser-Glu-His catalytic triad was found in five of six Ldc orthologs examined here, including FTL1678 and FTT0101. However, *C. jejuni* Pgp2, which has been shown to exhibit LdcA activity (Frirdich, 2014), lacks the Ser-Glu-His catalytic triad, indicating that a Ser-Glu-His catalytic triad may not be required for LdcA function. As such, we speculated that single amino acid mutations of the putative Ser134-Glu239-His308 catalytic triad in FTL1678 may not be sufficient to abolish enzyme function. Recombinant FTL1678 mutant proteins were generated and purified, each containing either two amino acid mutations (S134A/E239A, S134A/H308A, and E239A/H308A) or three amino acid mutations (S134A/E239A/H308A). Similar to what is described above, enzymatic assays were performed, using the TCT monomer as a substrate. While no specific activity to TCT monomer was detected for the empty vector control or buffer alone (Figure S2), WT FTL1678 was active against the TCT monomer (21.5 nmol/min/mg; Table S3). By comparison, no activity was detected for any of the double or triple mutant proteins (Table S3), indicating that mutation of two of more amino acids of the putative Ser134-Glu239-His308 catalytic triad ablates FTL1678 Ldc activity. Analysis of single amino acids of the putative Ser134-Glu239-His308 catalytic triad are described below.

### FTL1678 is OM-associated

*E. coli* and *P. aeruginosa* Ldcs have been localized to the bacterial cytoplasm (Templin, 1999, Korza, 2005). However, *C. jejuni* Pgp2 is unusual in that it contains a signal peptide and has been speculated to be periplasmic (Frirdich, 2014). In addition, *N. gonorrhoeae* LdcA was found to be periplasmic and outer membrane-associated (Lenz, 2017). As noted above, we previously demonstrated that FTL1678 is a DsbA substrate (Ren, 2014), indicating that FTL1678 is located in the *F. tularensis* envelope (*i.e.,* in the inner membrane [IM], periplasm, or outer membrane [OM]). Bioinformatic analyses of FTL1678 indicated that it is a periplasmic protein due to the presence of a signal peptide but absence of OM or lipoprotein signatures (Table S4). To experimentally confirm FTL1678 localization, we generated an *F. tularensis* strain with 6× histidine-tagged FTL1678, then performed spheroplasting, osmotic lysis, and sucrose density gradient centrifugation to separate IM and OM fractions and probe for protein subcellular localization. Immunoblotting of whole-cell lysates (WCL), OM fractions, and IM fractions demonstrated that the OM control protein, FopA (Huntley, 2007), only was present in WCL and OM fractions (but not IM fractions; Figure 2) and the IM control protein, SecY (Huntley, 2007), only was present in WCL and IM fractions (but not OM fractions; Figure 2). By comparison, FTL1678 only was detected in WCL and OM fractions, demonstrating OM-association (Figure 2). As noted above, because *N. gonorrhoeae* LdcA was found to fractionate to both the OM and soluble fractions (indicating periplasmic localization) (Lenz, 2017), we next examined the localization of *F. tularensis* periplasmic proteins in our fractions to better understand FTL1678 localization. TolB is a well-known periplasmic protein in Gram-negative bacteria and binds to PG due to its interaction with the peptidoglycan associated lipoprotein, Pal (Clavel, 1998, Walburger, 2002). We previously demonstrated that the *F. tularensis* Pal homolog is OM-localized (Huntley, 2007), similar to its OM-localization in other Gram-negative bacteria. Here, the *F. tularensis* TolB homolog was detected in OM fractions, but not IM fractions (Figure S4), indicating that *F. tularensis* PG-associated proteins also fractionate with OMs. In summary, data that FTL1678 is a DsbA substrate (*i.e.,* FTL1678 is an envelope protein)(Ren, 2014), FTL1678 contains a signal peptidase I cleavage site (Table S4), FTL1678 does not contain membrane protein signatures (Table S4), and FTL1678 is OM-associated (Figure 2), provides strong evidence that FTL1678 is not a cytoplasmic protein, unlike *E. coli* LdcA. Instead, our data indicate that, similar to *N. gonorrhoeae* LdcA, FTL1678 may be a periplasmic protein.

**Figure 2.**
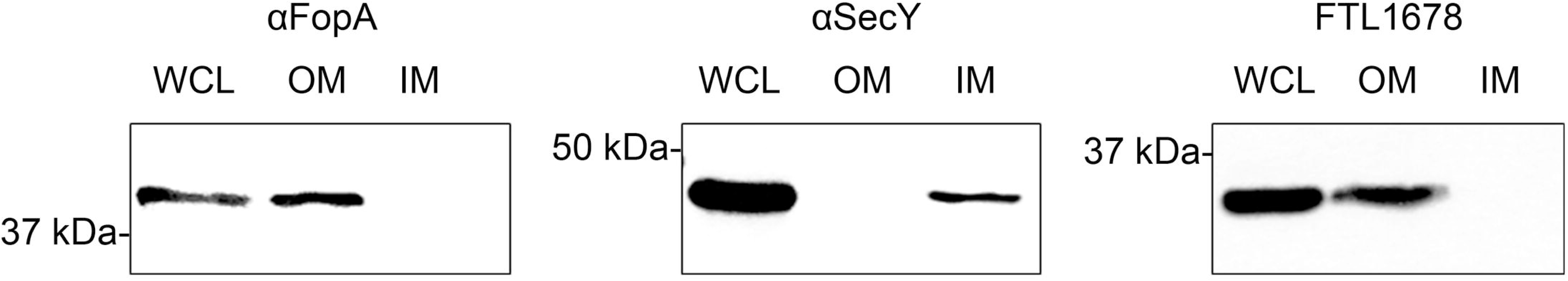
FTL1678 is OM-associated. Spheroplasting, osmotic lysis, and sucrose density gradient centrifugation were performed to separate inner membranes (IM) and outer membranes (OM) from *F*. *tularensis ΔFTL1678 trans*-complemented with a 6×histidine-tagged FTL1678. Whole-cell lysates (WCL), OM fractions, and IM fractions were separated by SDS-PAGE, transferred to nitrocellulose, and immunoblotting was performed using antisera specific for the OM control protein FopA (α FopA), IM control protein SecY (αSecY), or histidine-tagged FTL1678.

### Deletion of FTL1678 alters bacterial morphology

LdcA has been shown to be important for maintenance of bacterial morphology and structural integrity (Sycuro, 2013, Frirdich, 2014). In addition, mutations/deletions or combinations of mutations/deletions in PG-modifying proteins can result in abnormal bacterial morphology, emphasizing the importance of PG modification and recycling (Nelson, 2000, Guinane, 2006, Priyadarshini, 2007, Heidrich, 2001, Sycuro, 2010, Juan, 2018). To assess if FTL1678 plays a similar role in *F. tularensis*, we generated an isogenic deletion of FTL1678, referred to hereafter as Δ*FTL1678*, Δ*FTL1678*, in *F. tularensis* LVS. When examined by transmission electron microscopy (TEM), wild-type (WT) bacterial width ranged from 350 to 800 nm (Figures 3A and 3D), whereas Δ*FTL1678* bacteria were more uniform in cell width, averaging approx. 350 nm (Figures 3B and 3D). While WT bacteria generally were observed to be coccobacilli (Figure 3A), Δ*FTL1678* bacteria were found to be more coccoid in appearance, the Δ*FTL1678* OM was more tightly-associated than WT OMs, and three electron dense structures, likely the OM, PG, and IM, were present around the periphery of the majority of Δ*FTL1678* bacterium (Figure 3B), compared to less prominent outer structures surrounding WT bacteria (Figure 3A). Additionally, Δ*FTL1678* bacteria appeared more electron dense and had significantly-thicker OMs than WT bacteria (Figures 3A, 3B, and 3C).

**Figure 3.**
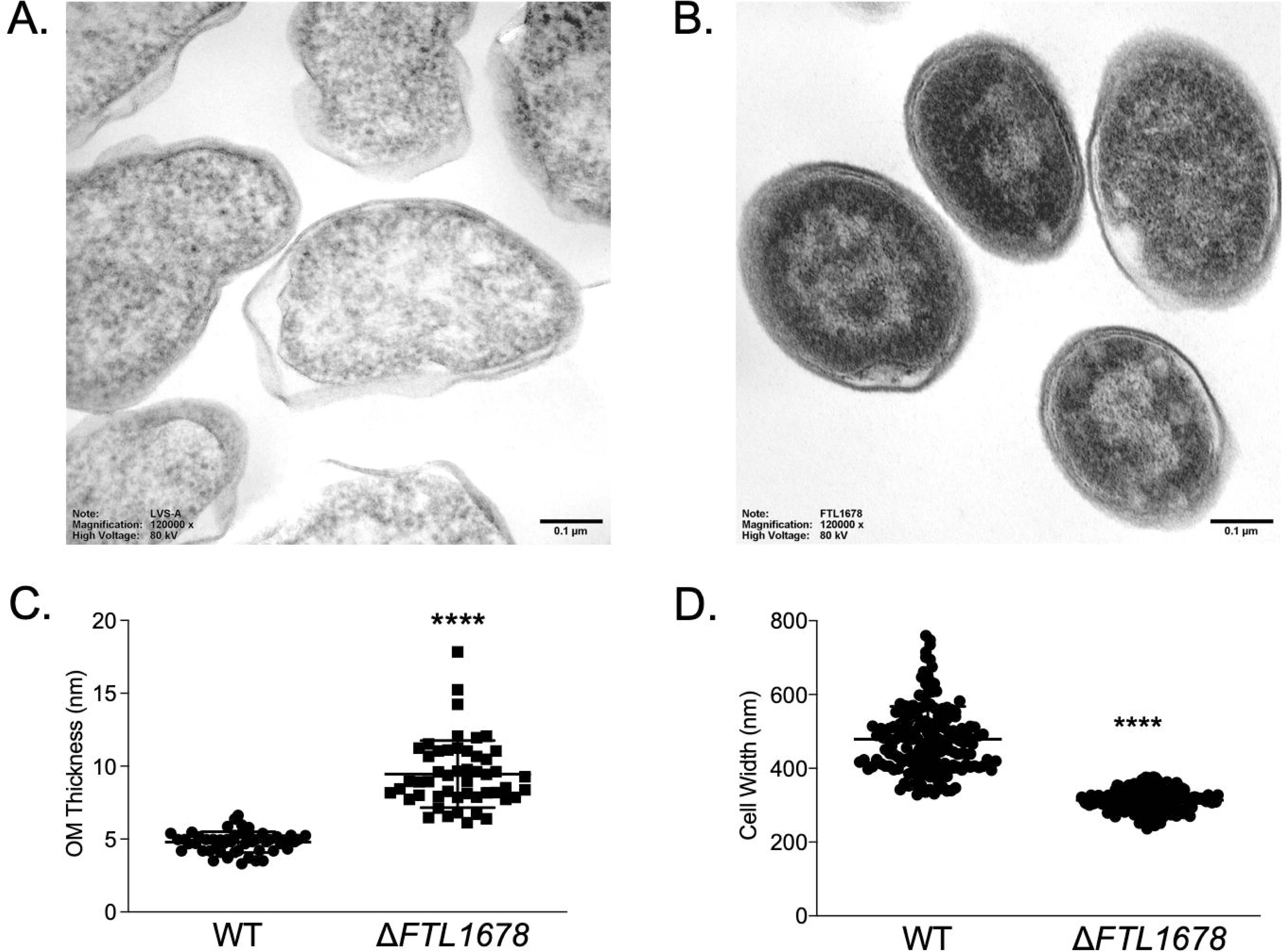
Deletion of FTL1678 alters bacterial morphology. Electron micrograph images of: (A) Wild-type LVS or (B) Δ*FTL1678* grown in sMHB to OD_600_ of 0.4. Scale bars represent 100 nm. Representative images shown; (C) Outer membrane [OM] thickness measurements [nm] were determined for WT and Δ*FTL1678*, with n=50 bacterial cells analyzed per experimental group, multiple width measurements recorded per individual bacterium, and average width/bacterium recorded; (D) Cell width measurements [nm] for WT and Δ*FTL1678*, with n=175 bacterial cells analyzed per experimental group. *\*\*\*\** indicates *P<*0.0001.

Previous studies have shown that deletion of genes for PG-modifying proteins (*e.g.,* murein hydrolases) can result in abnormal growth characteristics, including lysis during stationary phase (Templin, 1999) and an inability to separate daughter cells at the septa during cell division, resulting in abnormal bacterial chains (Heidrich, 2002, Denome, 1999, Priyadarshini, 2006, Priyadarshini, 2007, Heidrich, 2001, Chaput, 2016, Juan, 2018, Weaver *et al*., 2019). Although *N*-acetylmuramyl-L-alanine amidases have been shown to be predominantly involved in the cleavage of bacterial septa, deletion of lytic transglycosylases and some endopeptidases, in combination with amidase deletions, also have resulted in abnormal bacterial chains (Heidrich, 2001, Heidrich, 2002). To examine any potential replication defects of Δ*FTL1678*, we compared both OD_600_ values (Figure S5A) and CFUs over time (Figure S5B) of WT and Δ*FTL1678* in supplemented Mueller-Hinton Broth (sMHB; standard growth medium for *F. tularensis*; (Huntley, 2007)), finding that Δ*FTL1678* did not have any inherent growth defects. When examining both WT and Δ*FTL1678* by TEM for any septation defects or abnormal bacterial chains, approximately 10% of Δ*FTL1678* bacteria grew in chains of three to four bacteria (Figure S6A and S6B), whereas no WT bacteria exhibited this septation defect (data not shown). Taken together, our findings that Δ*FTL1678* is 1.5- to 2-times smaller than WT (Figure 3D), Δ*FTL1678* is more coccoid in shape (Figure 3B), Δ*FTL1678* has a thicker OM (Figure 3B and 3C), and Δ*FTL1678* has a partial septum defect (Figure S6), further support the role of FTL1678 as a PG-modifying enzyme that is important for bacterial elongation and division.

To provide additional evidence that alterations in Δ*FTL1678* morphology were solely due to loss of FTL1678 Ldc activity, we sought to complement Δ*FTL1678* with either FTL1678 or a known LdcA and assess restoration of WT bacterial morphology. Although *E. coli* and *P. aeruginosa* LdcA orthologs contain the Ser-Glu-His catalytic triad (Figure 1), those LdcA orthologs are cytoplasmic and, given our data that FTL1678 is OM-localized and PG-associated (*i.e.,* may be periplasmic; Figure 2), we speculated that cytoplasmic LdcA orthologs may not function in the *F. tularensis* periplasm. In contrast, *C. jejuni* Pgp2 has been shown to exhibit LdcA activity, has been speculated to be periplasmic (Frirdich, 2014), but lacks the Ser-Glu-His catalytic triad (Figure 1). To examine if *FTL1678* or *pgp2* could complement Δ*FTL1678*, we independently generated an *FTL1678 in trans*-complement and a *pgp2 in trans*-complement, examined bacterial morphologies of both complemented strains by TEM, and found that both complemented strains had similar morphologies as WT LVS (Figure S7A-B and Figure 3A). Additionally, OM thickness and cell width were measured for WT, Δ*FTL1678*, and both complemented strains, demonstrating that both complemented strains had OM thicknesses and cell widths similar to WT, and both complemented strains were significantly different from Δ*FTL1678* (Figure S7C-D). Taken together, these complementation studies provide further evidence that FTL1678 is an LdcA and that Δ*FTL1678* morphological changes are due to loss of LdcA activity. These studies also indicate that the *C. jejuni* LdcA ortholog, Pgp2, which is a putative periplasmic protein and lacks the Ser-Glu-His catalytic triad, exhibits LdcA activity in *F. tularensis*.

### Deletion of FTL1678 affects sensitivity to antibiotics, detergents, and stressors

Given the above noted morphological differences in the Δ*FTL1678* envelope (*e.g.*, thicker OM; tightly-associated OM; more electron dense envelope structures) using TEM, we assessed envelope (*i.e.,* IM, PG, and OM) integrity by growing both WT and Δ*FTL1678* bacteria in the presence of various antibiotics, detergents, and dyes, and measuring zones of inhibition after 48 h of growth (Table 2). Δ*FTL1678* was found to be more susceptible than WT to ampicillin, vancomycin, lysozyme, and SDS (Table 2), indicating potential changes to PG (ampicillin sensitivity), OM integrity (vancomycin and lysozyme sensitivity), or efflux pumps (SDS sensitivity). Conversely, Δ*FTL1678* was found to be more resistant than WT to gentamicin, tetracycline, chloramphenicol, ciprofloxacin, and ethidium bromide (Table 2). Given that the majority of these latter reagents must enter the cytoplasm to exert their inhibitory effects (*i.e.,* gentamicin, tetracycline, and chloramphenicol inhibit protein synthesis; ciprofloxacin and ethidium bromide interfere with DNA replication), these results suggest that Δ*FTL1678* bacteria exclude these inhibitory molecules from entering the cytoplasm.

To better understand potential differences in the Δ*FTL1678* envelope, WT and Δ*FTL1678* were grown in either sMHB at 37°C or in sMHB with various stress conditions. In sMHB at 37°C, Δ*FTL1678* did not exhibit a growth defect but, instead, appeared to grow to a higher optical density (OD_600_) than WT (Figure 4A). However, as noted above, despite higher OD_600_ measurements for Δ*FTL1678* at several time points, bacterial numbers were not significantly different between WT and Δ*FTL1678* (Figure S5). Although speculative, the disassociation between Δ*FTL1678* optical densities and bacterial numbers may be due to the observed TEM morphological differences of Δ*FTL1678* (Figure 3 and Figure S6). Compared with growth in sMHB at 37°C, no substantial differences in the growth rates of WT and Δ*FTL1678* were observed at either 40°C (Figure 4B) or in the presence of 60 μM CuCl_2_, an oxidizing agent (Ren, 2014); Figure 4C). However, Δ*FTL1678* grew considerably better than WT in the presence of 5 mM H_2_O_2,_ 5% NaCl, and pH 5.5 (Figures 4D, 4E, and 4F, respectively), providing further evidence of modifications to the Δ*FTL1678* envelope. To confirm that Δ*FTL1678* envelope integrity differences were not due to polar effects, both the *FTL1678 trans*-complement and a *pgp2 trans*-complement were grown in the presence of the same stressors and both complemented strains were found to exhibit similar phenotypes as WT LVS (Figure S8A-D and Table S5). Taken together, these results indicated that loss of *FTL1678,* and its associated LdcA activity, resulted in unidentified, and likely complex, perturbations in bacterial envelope components, including the OM, PG, and/or IM.

**Figure 4.**
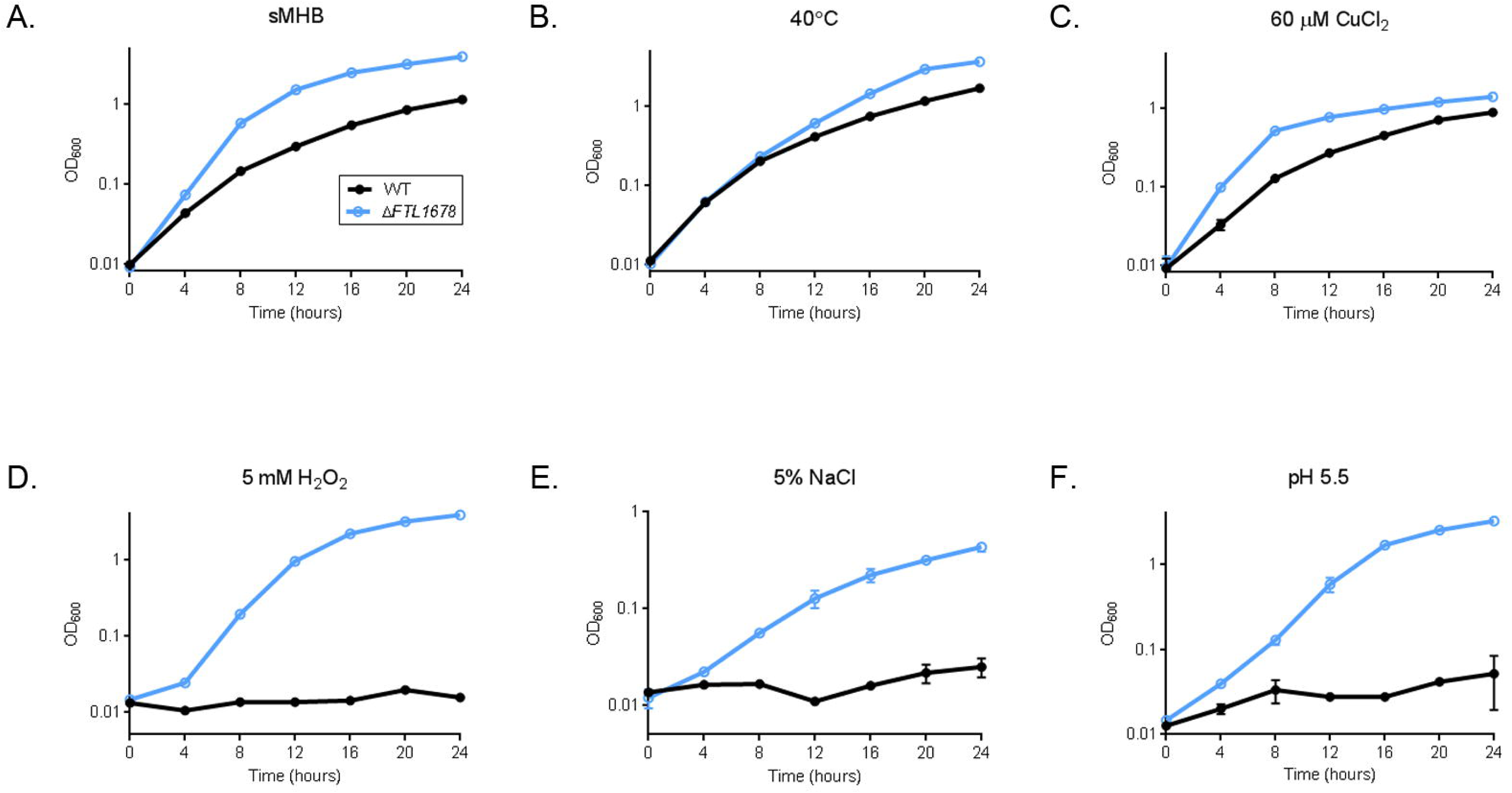
Deletion of *FTL1678* affects sensitivity to envelope stress. WT and Δ*FTL1678* were grown in 100 ml sMHB: (A) at 37°C, (B) at 40°C, (C) addition of 60 µM CuCl_2_, (D) addition of 5 mM H_2_O_2_, (E) addition of 5% NaCl, or (F) pH 5.5. Bacteria were grown in triplicate for 24 h and OD_600_ measurements were recorded every 4 h. Error bars represent standard deviation at each time point.

The Type A *F. tularensis* strain SchuS4 originally was isolated from a human tularemia patient and requires BSL3 containment. Given its relevance to human disease, we next generated an isogenic deletion mutant of the *FTL1678* homolog, *FTT0101*, in SchuS4. The susceptibilities of Δ*FTT0101* and WT SchuS4 to various antibiotics, detergents, and dyes were compared, with no significant differences observed (Table S6). At this time, we cannot fully explain why the Δ*FTL1678* mutant (*F. tularensis* Type B) displays altered sensitivity/resistance to antibiotics, detergents, and dyes (Table 2), while the Δ*FTL1678* mutant (*F. tularensis* Type A) did not demonstrate altered sensitivity/resistance toward these same compounds (compared to WT SchuS4; Table S6). However, this finding is not unexpected given genomic studies indicating that, despite >97% nucleotide identity between Type A and Type B *F. tularensis*, there are over 100 genomic rearrangements between Type A and Type B and each subspecies encodes over 100 unique genes that likely influence known differences in Type A and Type B virulence (Petrosino *et al*., 2006).

### FTL1678 and LdcA activity are required for F. tularensis virulence in vivo

To examine if FTL1678 plays a role in *F. tularensis* virulence, groups of C3H/HeN mice were intranasally infected with 10^4^ CFU of either WT or Δ*FTL1678* and monitored daily for signs of disease. Whereas all WT-infected mice died by day 9 post-infection (median time-to-death day 6), Δ*FTL1678* was completely attenuated (100% survival through day 21 post-infection), demonstrating that FTL1678 is required for *F. tularensis* virulence (Figure 5A). To confirm that the observed attenuation was solely due to the deletion of *FTL1678*, and not to polar effects, we tested the virulence of the Δ*FTL1678 in trans*-complement in mice, which fully-restored virulence to WT levels (all mice died by day 7; median time-to-death day 6; Figure 5A).

**Figure 5.**
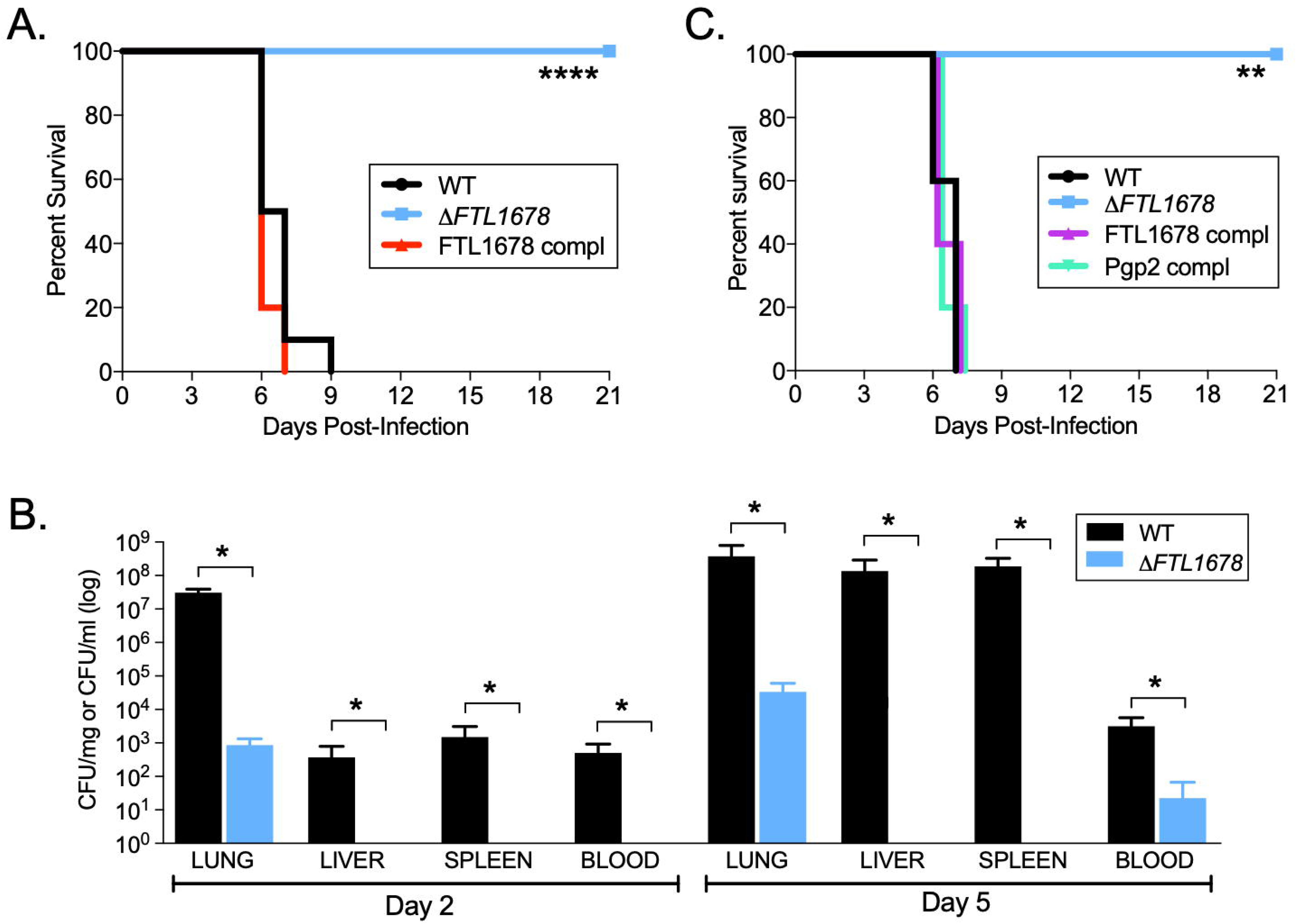
Δ*FTL1678* is fully-attenuated in a mouse pulmonary infection model. (A) Groups of 5 C3H/HeN mice were intranasally-infected with 10^5^ CFU of either wild-type WT, Δ*FTL1678*, or Δ*FTL1678 trans*-complemented with Δ*FTL1678* [FTL1678 compl]. Animal health was monitored daily through day 21 post-infection. **** indicates *P*<0.0001; (B) Lungs, livers, spleens, and blood were aseptically harvested from mice infected with 10^4^ CFU of either WT or Δ*FTL1678* on days 2 and 5 post-infection and plated to enumerate bacterial numbers. * indicates *P*<0.01; (C) Groups of 5 C3H/HeN mice were intranasally-infected with 10^5^ CFU of either LVS, Δ*FTL1678*, FTL1678 *trans*-complement [FTL1678 compl], or *C. jejuni* Pgp2 *trans*-complement [Pgp2 compl]. Animal health was monitored through day 21 post-infection. ** indicates *P*<0.01.

To more carefully assess Δ*FTL1678* attenuation *in vivo*, we intranasally-infected mice with 10^4^ CFU of either WT LVS or Δ*FTL1678*, and enumerated bacterial CFUs from lungs, livers, spleens, and blood on days 2 and 5 post-infection to examine bacterial replication and dissemination to these organs/tissues over time. On day 2 post-infection, WT LVS replicated to >10^7^ CFU/mg lung and had disseminated to livers, spleens (approx. 10^3^ CFU/mg), and blood (10^3^ CFU/ml; Figure 5B). In contrast, Δ*FTL1678* had an initial (day 2) colonization defect in the lungs (>35,000-fold lower than WT) and was unable to disseminate to livers, spleens, or blood (Figure 5B). By day 5 post-infection, the attenuation of Δ*FTL1678* was even more apparent, with WT LVS replicating to extremely high numbers (approx. 10^8^ CFU/mg) in lungs, livers, and spleens, compared with Δ*FTL1678,* which replicated in lungs between day 2 and 5, but was >11,000-fold attenuated in lungs and was not detectable in livers or spleens (Figure 5B). Although Δ*FTL1678* was detected in the blood on day 5, it was 142-fold less than WT LVS (Figure 5B).

As noted above, of the six Ldc orthologs examined here, only *C. jejuni* Pgp2 lacks the Ser-Glu-His catalytic triad (Figure 1). However, *C. jejuni* Pgp2 has been shown to exhibit LdcA activity (Frirdich, 2014), indicating that a Ser-Glu-His catalytic triad is not required for LdcA function. To test if an LdcA ortholog, without the Ser-Glu-His catalytic triad, could restore virulence in the Δ*FTL1678* mutant, we complemented Δ*FTL1678* with *C. jejuni* Pgp2 *in trans* and infected groups of mice with either WT LVS, Δ*FTL1678*, Δ*FTL1678*, *FTL1678 trans*-complemented with FTL1678, or Δ*FTL1678 trans*-complemented with *C. jejuni* Pgp2. While Δ*FTL1678* was fully attenuated (100% survival through day 21), the Pgp2 *trans*-complement was fully-virulent (median time-to-death 6 days; all mice dead by day 7), nearly identical to WT LVS (median time-to-death 7 days; all mice dead by day 7) and the FTL1678 *trans*-complement (median time-to-death 6 days; all mice dead by day7; Figure 5C). These *in vivo* data provide further evidence that FTL1678 is an Ldc and that Ldc activity, with or without the Ser-Glu-His catalytic triad, is required for *F. tularensis* virulence.

Finally, given the relevance of SchuS4 to human disease, we next examined the virulence of Δ*FTT0101*, the *FTL1678* homolog, in our mouse infection model. When groups of C3H/HeN mice were intranasally-infected with either WT SchuS4 or Δ*FTL1678*, all mice died by day 7 post-infection, indicating that *FTT0101* is not required for SchuS4 virulence (Figure S9). Whereas Δ*FTT0101*-infected mice exhibited a slightly delayed time-to-death (median time-to-death day 6; Figure S9), compared with WT SchuS4-infected mice (median time-to-death day 5; Figure S9), this may be due to differences in the infectious dose administered to mice in this experiment (80 CFU SchuS4; 12 CFU Δ*FTT0101*). However, it also remains possible that the extreme virulence of SchuS4 (intranasal LD_50_ <10 CFU in our hands) and over 100 genomic rearrangements between Type A and Type B *F. tularensis* (noted above) complicates assessments of mutant attenuation *in vivo*.

### Individual residues of the FTL1678 catalytic triad are not required for F. tularensis virulence in vivo

As noted above and highlighted in Figure 1, *P. aeruginosa* LdcA contains a Ser-Glu-His catalytic triad which is essential for function and is characteristic of Ldc in the Peptidase_S66 family (Korza, 2005). The same catalytic triad also has been confirmed in Ldc from *E. coli* (Meyer, 2018), *Novosphingobium aromaticivorans* (Das, 2013), and *N. gonorrhoeae* (Lenz, 2017). Given the relatively conserved spacing of Ser134-Glu239-His308 residues in FTL1678 (Figure 1) and our findings that both double (S134A/E239A, S134A/H308A, E239A/H308A) and triple (S134A/E239A/H308A) mutants did not exhibit LdcA activity (Table S3), we tested if individual amino acid residues of the catalytic triad were required for *F. tularensis* virulence (similar to Figure 5A virulence assessments for Δ*FTL1678* and the FTL1678 complemented strain). Site-directed mutagenesis was performed to independently generate FTL1678 complementation constructs containing either S134A, E239A, or H308A mutations. Next, Δ*FTL1678* was complemented *in*-*trans* with each of these FTL1678 catalytic triad point mutants, and groups of C3H/HeN mice were intranasally infected with either WT, Δ*FTL1678*, *FTL1678 trans*-complemented with one of the FTL1678 catalytic triad point mutants (referred to hereafter as S134A, E239A, and H308A). Confirming our previous findings, Δ*FTL1678* was completely attenuated (100% survival through day 21), while complementation of Δ*FTL1678* with either FTL1678 (all mice dead by day 8), S134A (all mice dead by day 7), E239A (all mice dead by day 8), or H308A (all mice dead by day 8), fully-restored virulence to WT LVS levels (all mice dead by day 10; Figure S10). These results should not be overinterpreted, as it is difficult to directly compare or fully explain why mutations of single amino acids in the FTL1678 putative catalytic triad had no effect on *in vivo* virulence (Figure S10), while mutations of any two amino acids in the catalytic triad ablated FTL1678 LdcA enzyme activity *in vitro* (Table S3). It remains possible that single or multiple residues of the FTL1678 catalytic triad are required for full LdcA activity *in vitro* or that, in the context of the whole bacterium (*i.e., in vivo*), other *F. tularensis* proteins may have compensated for partially-functional FTL1678, due to S134A, E239A, and H308A single amino acid mutations. Regardless, these studies indicate that, while single amino acid mutations of the FTL1678 Ser-Glu-His catalytic triad do not impact *F. tularensis* virulence *in vivo*, two or more residues of the Ser-Glu-His catalytic triad are required for *FTL1678* LdcA activity.

### FTL1678 is required for F. tularensis replication in macrophages

*F. tularensis* is an intracellular pathogen and macrophages appear to be one of the major targets for *F. tularensis* infection and replication (De Pascalis, 2018, Steiner, 2017, Hall, 2008). To investigate potential replication defects of Δ*FTL1678* in macrophages, J774A.1 macrophages or murine bone marrow-derived macrophages (mBMDM) were infected with either WT LVS or *FTL1678* (MOI 100:1) and bacterial numbers were enumerated at 0 h (entry), 6 h, and 24 h post-infection. At entry (0 h), > 300-fold more Δ*FTL1678* were present in both macrophage lines, compared with WT LVS (Figure 6A). This likely was due to the above noted gentamicin resistance of Δ*FTL1678* (Table 2). Attempts to normalize entry numbers for both WT LVS and Δ*FTL1678*, using different antibiotics or combinations of antibiotics, were not successful. Despite higher numbers of Δ*FTL1678* in both macrophages at entry (0 h) and 6 h, Δ*FTL1678* was unable to replicate in either macrophage, decreasing 13-fold in BMDM and 5-fold in J774A.1 macrophages from 6 h to 24 h (CFU data in Figure 6A; fold change data in Figure 6B). By comparison, WT LVS numbers increased 236-fold in BMDM and 22-fold in J774A.1 macrophages from 6 h to 24 h (Figure 6A-B). Taken together, these *in vitro* results (Figure 6A-B) confirm the observed *in vivo* attenuation of Δ*FTL1678* (Figure 5A-C).

**Figure 6.**
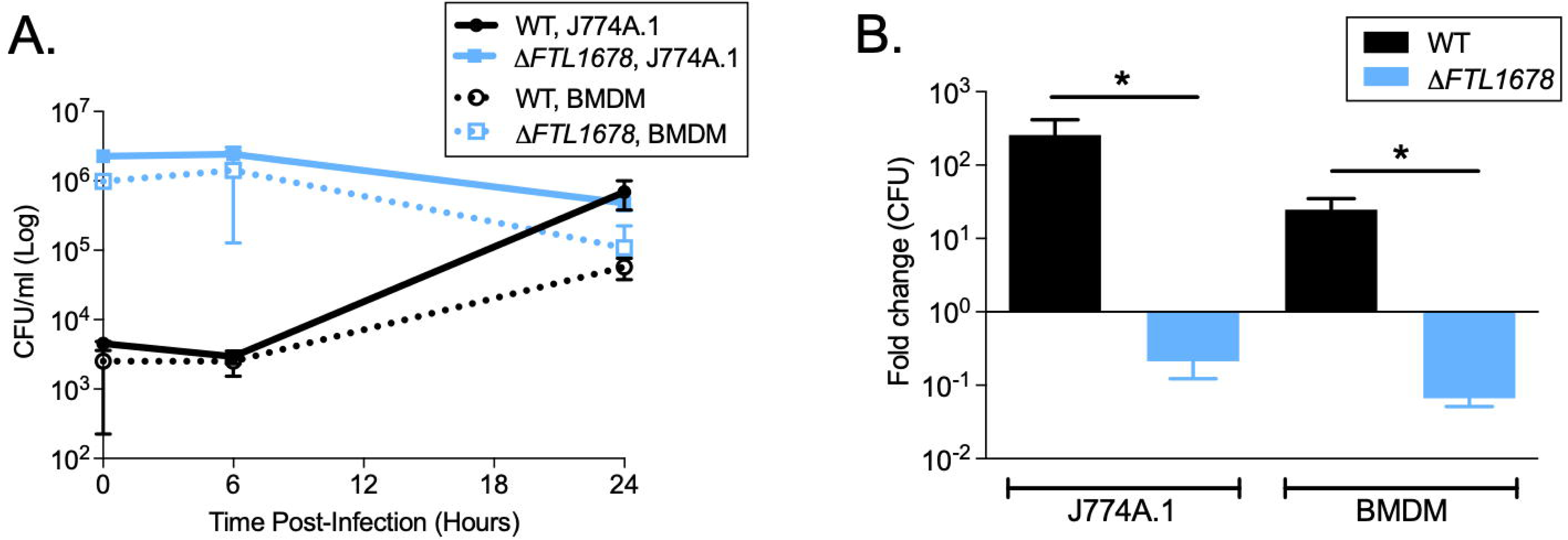
FTL1678 is required for *F. tularensis* replication in macrophages. (A) J774A.1 macrophages or mouse bone marrow-derived macrophages (mBMDMs) were infected with WT or Δ*FTL1678* at an MOI of 100:1 and bacterial numbers were enumerated at entry (0 h), 6 h, and 24 h post-infection. (B) Fold change in bacterial numbers from 6 to 24 h post-infection was calculated. * indicates *P*< 0.01.

### ΔFTL1678 protects mice against Type A F. tularensis infection

No FDA-approved vaccine currently is available to prevent tularemia. In addition, *F. tularensis* is designated as an NIH Category A priority pathogen and CDC Tier 1 Select Agent, highlighting the extreme virulence of this bacterium and the need for a safe and effective vaccine to prevent tularemia. Given our above findings that 10^5^ CFU of Δ*FTL1678* did not cause disease or death in mice (Figure 5A-C), we next examined whether high doses (10^7^ or 10^9^ CFU) of Δ*FTL1678* were attenuated or if Δ*FTL1678* immunization could protect mice from fully-virulent Type A *F. tularensis* SchuS4 challenge. First, all mice intranasally immunized with either 10^5^, 10^7^, or 10^9^ CFU of Δ*FTL1678* survived through day 28 post-infection, with no signs of clinical disease (Figure 7A). Next, on day 29, all mice were boosted with 10^9^ CFU of Δ*FTL1678* and no mice demonstrated any signs of disease through day 50 (Figure 7A). Finally, on day 51, mice were intranasally-challenged with 120 CFU (6× the LD_50_) of SchuS4 and the health status of each immunization group was monitored for 26 days post-challenge. In a dose-dependent manner, the 10^9^ prime-10^9^ boost regimen conferred 80% protection, the 10^7^ prime-10^9^ boost regimen conferred 40% protection, and the 10^5^ prime-10^9^ boost regimen conferred 20% protection (Figure 7B). These data demonstrate that Δ*FTL1678* is highly attenuated (up to 10^9^ CFU) and that Δ*FTL1678* may be able to be used as a live, attenuated vaccine.

**Figure 7.**
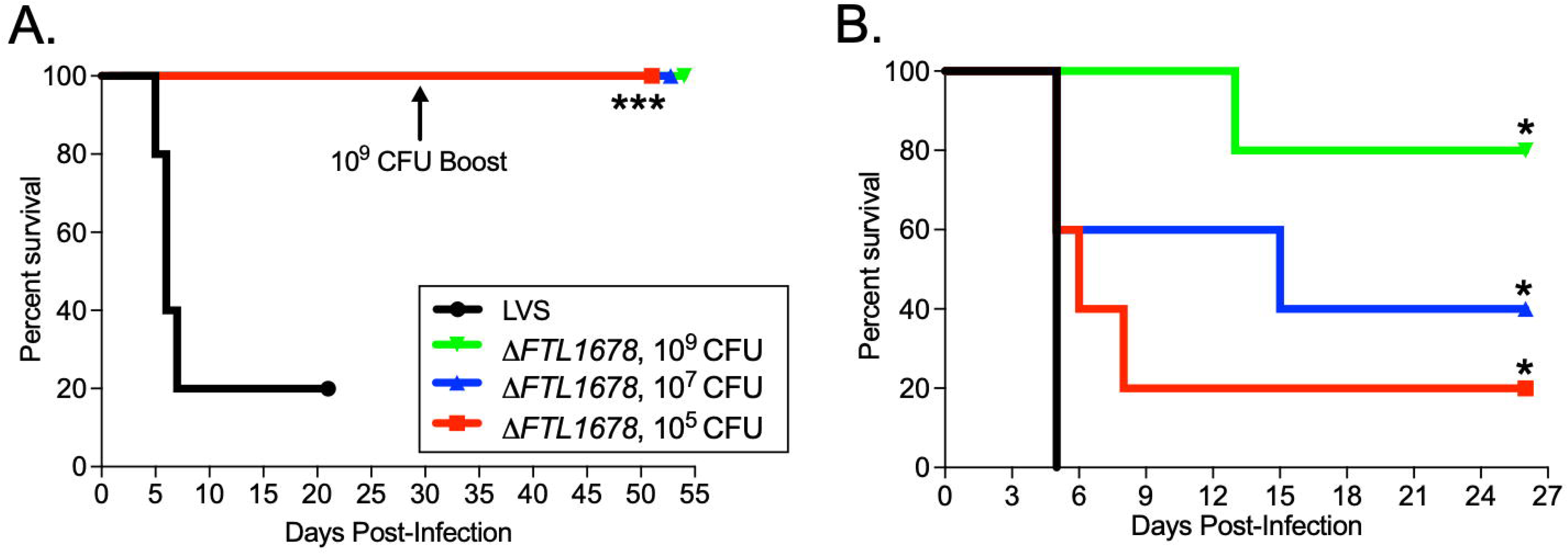
Δ*FTL1678* protects against fully-virulent Type A *F. tularensis* SchuS4. (A) Groups of 5 C3H/HeN mice were intranasally infected with either 10^5^ CFU WT or 10^5^, 10^7^, or 10^9^ CFU Δ*FTL1678*. On day 29 post-infection, mice were boosted with 10^9^ CFU Δ*FTL1678* and animal health was monitored daily through day 50 post-infection. *** *P*<0.001; (B) Mice from A were intranasally-challenged with 120 CFU of wild-type SchuS4 [BSL3; 6× LD50]. Animal health was monitored daily through day 26 post-infection. * indicates *P*<0.001.

### ΔFTL1678 does not cause tissue damage

The *in vitro* (Figure 6) and *in vivo* (Figure 5A-C and Figure 7A) attenuation of Δ*FTL1678,* as well as protection against SchuS4 pulmonary challenge (Figure 7B), indicated that Δ*FTL1678* could be used as a live, attenuated vaccine. While live, attenuated vaccines have been extremely effective at preventing a number of diseases, they can pose safety challenges (Minor, 2015, Roberts, 2018). To assess whether Δ*FTL1678* immunization induced any pathology in immunized mice, lungs, livers, and spleens from uninfected, WT LVS-, or Δ*FTL1678*-infected mice were assessed for pathologic changes on day 5 post-infection/immunization. Day 5 is when mice exhibit severe signs of disease and is one day before the majority of WT-infected mice begin succumbing to disease (Figures 5 and 7). WT LVS-infected lungs demonstrated alveolar wall thickening, large areas of inflammation, and severe neutrophil infiltration (Figure 8A). By comparison, little inflammation was observed in Δ*FTL1678*-infected lungs, although some red blood cell congestion was present, indicating a limited, acute immune response that was quickly resolved (Figure 8A). Whereas WT LVS-infected livers were characterized by diffuse inflammation with focal areas of necrosis, Δ*FTL1678*-infected livers were virtually indistinguishable from uninfected livers, with no observable pathology (Figure 8A). Finally, although the architecture of WT LVS-infected spleens lacked distinct areas of white pulp or red pulp, indicative of a severe infection, Δ*FTL1678*-infected spleens were observed to contain distinct areas of red pulp and white pulp, with some red blood cell congestion – indicating a limited, acute immune response that was quickly resolved (Figure 8A). All tissues were blindly scored using a pathology severity index (scale from 0 to 4, with 4 indicating severe pathology), confirming that Δ*FTL1678*-infected tissues were virtually indistinguishable from uninfected tissues (pathology scores of 1 for lungs, 0 for liver, and 1.5 for spleens) and WT LVS-infected tissues had significantly higher pathology scores (pathology scores >3.5 for all tissues; Figure 8B).

**Figure 8.**
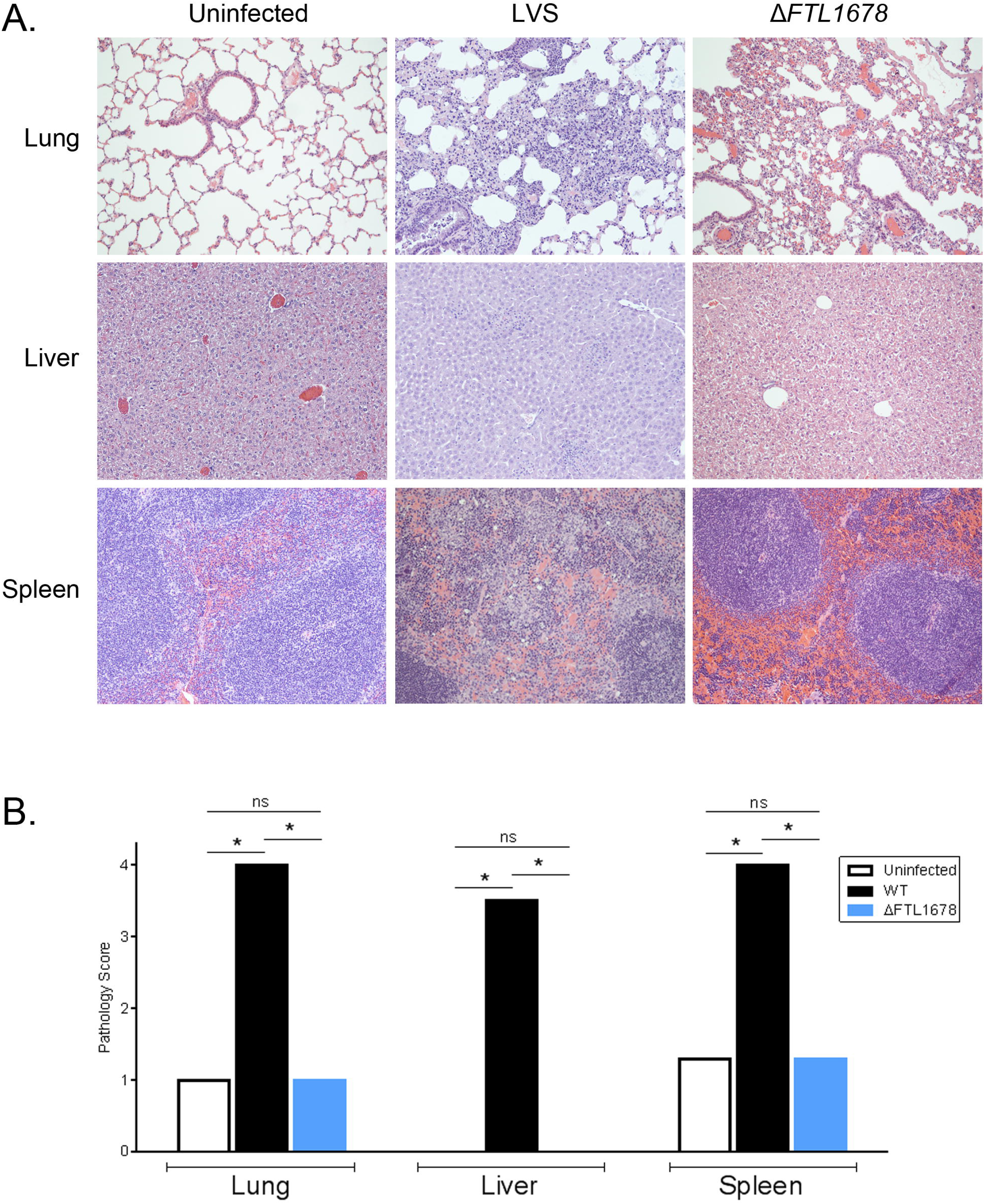
Δ*FTL1678* does not induce tissue damage. (A) Hematoxylin and eosin (H&E)-stained lungs, livers, and spleens were examined from either uninfected, *F. tularensis* WT LVS-, or Δ*FTL1678*-infected mice at 10× objective. (B) Tissues were graded on a scale of 0 to 4, with 4 being the most severe. * indicates *P*<0.05.

## Discussion

Bacterial PG is a complex, mesh-like structure, composed of a glycan backbone, crosslinked to varying degrees, by peptide chains (de Pedro, 2015). It is well known that this structure plays an important role in maintaining Gram-negative bacterial cell morphology, membrane integrity, regulating changes in osmotic pressure, and providing a platform for attachment of the OM (den Blaauwen, 2008, Silhavy *et al*., 2010). Although a majority of PG studies have focused on how the thick layer of PG in Gram-positive bacteria contributes to virulence and antibiotic resistance, more recent studies have highlighted that Gram-negative PG also is intimately linked with pathogenicity (Juan, 2018).

PG recycling is an essential function of Gram-negative bacteria during cell growth and division to produce new cell wall components. In fact, Gram-negative bacteria recycle up to 60% of their PG with every generation, suggesting that both PG synthesis and PG recycling are dynamic (Dhar, 2018, Typas, 2011, Park, 2008). A number of proteins are involved in these processes and, while they are well-characterized in *E. coli*, very little is known about these pathways in intracellular pathogens such as *Burkholderia pseudomallei*, *Legionella pneumophila*, or *F. tularensis* (van Heijenoort, 2011, Jenkins, 2019, Spidlova, 2018, Kijek, 2019).

*E. coli* LdcA, a cytoplasmic protein, was the first L,D-carboxypeptidase to be identified and was shown to be important for PG recycling and survival during stationary phase (Ursinus, 1992, Templin, 1999). More recently, Ldc orthologs have been identified in *P. aeruginosa* (Korza, 2005), *C. jejuni* (Frirdich, 2014), *N. gonorrhoeae* (Lenz *et al*., 2017), and *N. aromaticivorans* (Das, 2013). In this study, we identified an *F. tularensis* Ldc ortholog, FTL1678, which we propose naming LdcA based on its confirmed L,D, carboxypeptidase activity (Table 1) and role in maintaining bacterial morphology. Unlike well-characterized cytoplasmic LdcA orthologs from *E. coli* and *P. aeruginosa*, we demonstrated that *F. tularensis* LdcA was localized to OM fractions and, given co-localization with PG-associated proteins Pal and TolB, is most likely located on the inner leaflet of the OM or in the periplasm (associated with PG). At this time, we can only speculate on the OM-association or periplasmic localization of *F. tularensis* LdcA, but in the context of PG repair and recycling, periplasmic LdcA certainly offers a fitness advantage. In addition, this is not the first report of a periplasmic LdcA, as *C. jejuni* Pgp2 is predicted to be periplasmic and *N. gonorrhoeae* LdcA previously was reported to be periplasmic (Lenz, 2017, Frirdich, 2014).

Our results demonstrated, for the first time, that *F. tularensis* LdcA directly acts on the TCT tetrapeptide and the reducing PG monomer. More importantly, we demonstrated that *F. tularensis* LdcA directly cleaves PG pentapeptides to tripeptides, without a prior cleavage event by a D,D-carboxypeptidase/penicillin binding protein (PBP), such as DacD, and that FTL1678 cleaves TCT dimers with 4-3 and 3-3 cross links, highlighting that *F. tularensis* LdcA is a multi-functional enzyme that exhibits both L,D-carboxypeptidase and L,D-endopeptidase activities (Table 1 and Table S2). It should be noted that although *N. gonorrhoeae* LdcA also has been reported to have L,D-endopeptidase activity, this activity was shown to cleave tetra-tri and tri-tri dimers, but not tetra-tetra dimers, suggesting a specificity for 3-3 cross linked dimers (Lenz, 2017). Interestingly, only two previous studies have examined putative *F. tularensis* PG modifying enzymes and both studies primarily focused on the role of an *F. tularensis* DacD ortholog in virulence, with no PG activity assays to confirm function (Spidlova, 2018, Kijek, 2019). In our PG cleavage analysis, *F. tularensis* LdcA demonstrated the highest specific activity on disaccharide-tetrapeptide PG substrates (GlcNAc-anhydroMurNAc-L-Ala-γ-D-Glu-*meso*-A_2_pm-D-Ala [TCT] and GlcNAc-MurNAc-L-Ala-γ-D-Glu-*meso*-A_2_pm-D-Ala [reducing PG monomer]), followed by cleavage of pentapeptide PG substrates (MurNAc-L-Ala-γ-D-Glu-*meso*-A_2_pm-D-Ala-D-Ala and UDP-MurNAc-L-Ala-γ-D-Glu-*meso*-A_2_pm-D-Ala-D-Ala). Despite high specific activity of *F. tularensis* LdcA on tetrapeptide attached to the disaccharide, *tularensis* LdcA demonstrated approximately 6-times lower specific activity on free tetrapeptide (no sugars) (Table 1). In contrast, *E. coli* LdcA has been shown to have the highest specific activity on free tetrapeptide, monosaccharide-tetrapeptide (MurNAc-L-Ala-γ-D-Glu-*meso*-A_2_pm-D-Ala), and monosaccharide tetrapeptide linked to a glycan lipid carrier (UDP-MurNAc-tetrapeptide) (Templin, 1999), but is unable to cleave dimeric muropeptides. Additionally, *F. tularensis* LdcA was active against different forms of dimers (two TCT monomers carrying tri- or tetrapeptide chains connected either by a 4-3 or a 3-3-crosslink) but was not active on the PG polymer. The cleavage of the latter dimers indicated that *F. tularensis* LdcA possesses both L,D-endopeptidase and L,D,-carboxypeptidase activities and could cleave the A_2_pm-A_2_pm, A_2_pm-D-Ala and A_2_pm-Gly peptide bonds (L-D bonds in all cases) present in these dimers, with more or less efficacy.

Previous studies have shown that Ldc orthologs are important for bacterial morphology and membrane integrity. Deletion of the *ldc* orthologs *csd6* from *H. pylori* (Sycuro, 2013) and *pgp2* from *C. jejuni* (Frirdich, 2014) resulted in loss of helical morphology. Here, we demonstrated that FTL1678 is essential for maintaining both the size (width) and the coccobacillus morphology of *F. tularensis,* as Δ*FTL1678* bacteria were significantly-smaller than WT and exhibited a more-rounded, cocci shape than WT (Figure 3). Further evidence for the role of *F. tularensis* LdcA in modifying and recycling PG, which impacts bacterial morphology, is provided by TEM images of Δ*FTL1678* bacteria that have prominent three-layered structures at their periphery, including a thick middle layer (presumably PG), compared to WT (Figure 3). In Δ*FTL1678* bacteria, it is possible that loss of LdcA activity may have reduced PG recycling or may have affected the breakdown of existing PG (important for cell division), resulting in a buildup of pentapeptides or tetrapeptides that are highly-crosslinked. Our lab and others have repeatedly attempted to isolate and analyze *F. tularensis* PG but theses attempts have not been successful. As such, we can only speculate on the true nature of the thick PG and OM layers in Δ*FTL1678* (Figure 3).

Because Δ*FTL1678* bacteria were found to have a thicker OM (Figure 3C), a prominent middle layer in their envelope (presumably PG; Figure 3B), and an altered cell morphology (Figure 3B), we investigated differences in WT and Δ*FTL1678* susceptibility to various antibiotics, detergents, and stressors. Vancomycin and lysozyme, usually not effective against Gram-negative species due to their inability to penetrate the OM, inhibited Δ*FTL1678* growth (Table 2), indicating increased permeability of the Δ*FTL1678* OM. Vancomycin, in particular, may have been effective on Δ*FTL1678* because its mechanism of action includes binding to the two terminal D-Ala-D-Ala residues of PG pentapeptide chains and preventing cross-linking of monomers. Δ*FTL1678* may have increased amounts of pentapeptides present in its PG, providing more targets for vancomycin action. Similarly, ampicillin inhibits bacterial transpeptidases, which blocks cross-linking of peptide side chains of PG strands. Taken together, the enhanced susceptibility of Δ*FTL1678* bacteria to vancomycin and ampicillin supports the role of *F. tularensis* LdcA as a PG-modifying enzyme. Conversely, Δ*FTL1678* was more resistant to antibiotics and molecules that must cross the IM to exert their toxic effects, including gentamicin, tetracycline, chloramphenicol, ciprofloxacin, and ethidium bromide (Table 2), suggesting that Δ*FTL1678* bacteria have a less permeable IM. Indeed, decreased permeability of the IM and altered activity of IM efflux pumps may help explain the more electron-dense staining of Δ*FTL1678*, compared to WT (Figure 3). Many mechanisms may explain why Δ*FTL1678* bacteria were more resistant to other stressors, including H_2_O_2_, high salt, and low pH, including increased expression/activity of chaperone proteins, efflux pumps, antioxidant/scavenger proteins, and membrane stabilizing proteins (Mishra & Imlay, 2012, Knodler *et al*., 2003, Lund *et al*., 2014). Future studies are needed to define these mechanisms, which are likely to be complex.

The extreme virulence of Type A *F. tularensis* and its designation as a Tier 1 Select Agent highlight why studies to identify *F. tularensis* virulence factors and the development of new vaccines is important. In this study, we identified the role of a previously unstudied protein, FTL1678, in PG recycling, PG integrity, and bacterial morphology. In addition, we found that FTL1678 was required for *F. tularensis* LVS virulence and demonstrated that Δ*FTL1678* conferred 80% protection against fully-virulent, Type A *F. tularensis* SchuS4 pulmonary challenge. Further studies are needed to determine specific immune responses induced by Δ*FTL1678* immunization, as well as to identify the most effective immunization regimen (*e.g.,* number of immunizations and time between immunizations).

Finally, given our findings that PG maintenance and recycling are important for *F. tularensis* virulence, and that future studies may reveal additional PG-associated enzymes, we used bioinformatic approaches to predict other proteins involved in *F. tularensis* PG synthesis and recycling (Figure 9). While at least seven PG synthesis and recycling genes/proteins orthologs could not be identified, 22 putative PG synthesis and recycling proteins were identified in *F. tularensis* (Figure 9). Of these, only DacD (FTL1060/FTT1029) has been studied in *F. tularensis*. Given our observed attenuation of Δ*FTL1678*, future studies to better understand PG synthesis and recycling pathways may offer more opportunities to better understand the virulence of *F. tularensis* and other intracellular pathogens. Characterization of other proteins involved in PG pathways may provide clues as to why *F*. *tularensis* LdcA is OM-associated or periplasmic and encodes both L,D-carboxypeptidase and L,D-endopeptidase activities.

**Figure 9.**
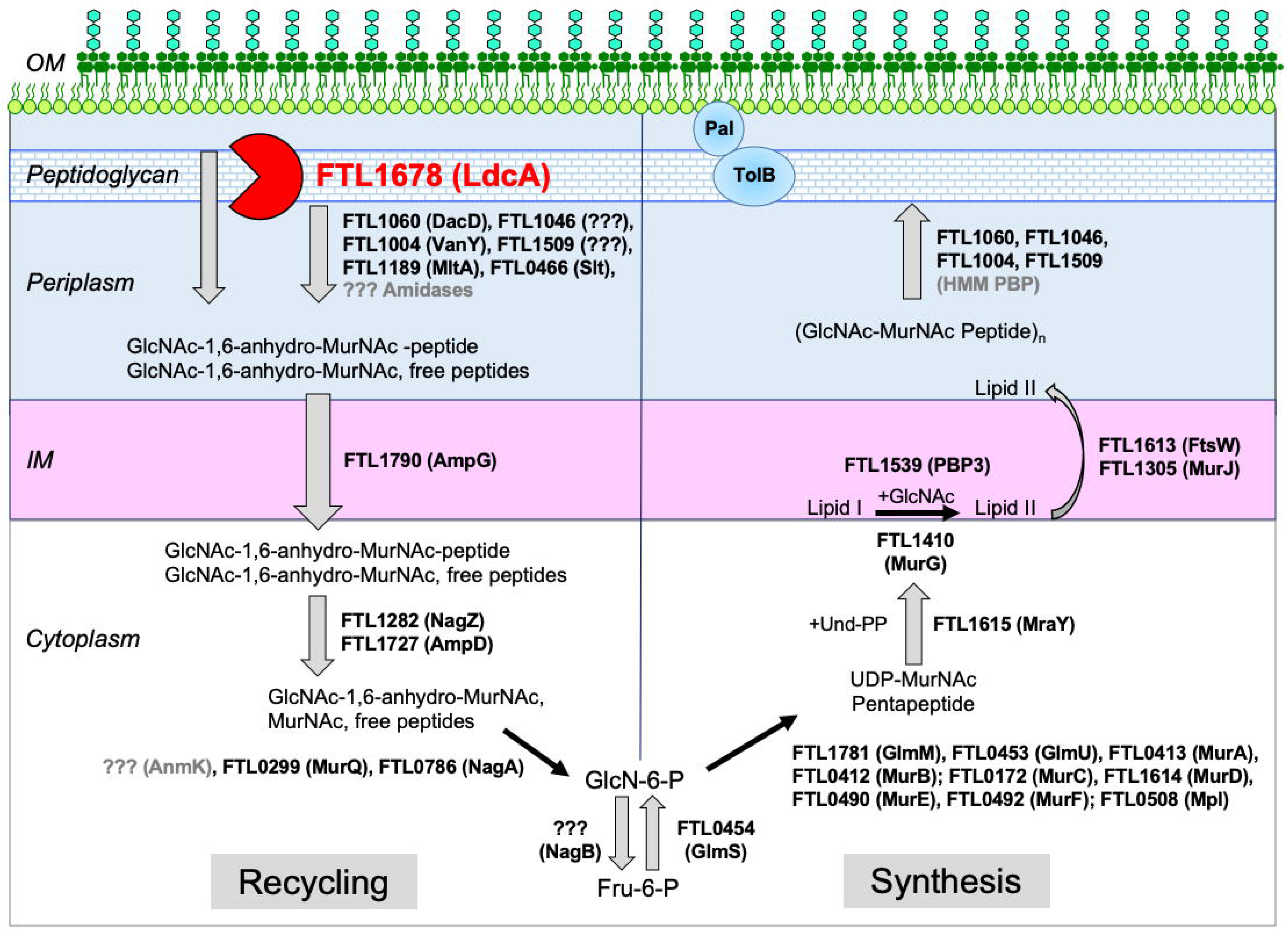
Model of *F. tularensis* peptidoglycan synthesis and recycling pathways. Bioinformatic analyses were used to predict proteins that may be involved in peptidoglycan synthesis and recycling in *F. tularensis*. *F. tularensis* LVS gene locus tags are indicated, with *E. coli* or Gram-negative ortholog protein names. OM, outer membrane. IM, inner membrane. GlcNAc, *N*-acetylglucosamine. MurNAc, *N*-acetylmuramic acid. Pal, OM-localized peptidoglycan-associated lipoprotein. TolB, periplasmic protein that interacts with Pal and peptidoglycan. HMM PBP, high molecular weight penicillin binding protein. LMM PBP, low molecular weight penicillin binding protein.

## Experimental Procedures

### Bacterial strains and culture conditions

*F. tularensis* Type A strain SchuS4 and *F. tularensis* Type B strain LVS were obtained from BEI Resources and cultured as previously described (Wu, 2016, Ren, 2014). All experiments with SchuS4 were performed under BSL3 containment conditions at the University of Toledo Health Science Campus BSL3 laboratory. Routine *F. tularensis* cultures were grown overnight at 37°C with 5% CO_2_ on supplemented Mueller-Hinton agar (sMHA): Mueller-Hinton broth powder (Becton Dickinson) was mixed with 1.6% (wt/vol) Bacto Agar (Becton Dickinson), autoclaved, and further supplemented with 2.5% (vol/vol) bovine calf serum (Hyclone), 2% (vol/vol) IsoVitaleX (Becton Dickinson), 0.1% (wt/vol) glucose, and 0.025% (wt/ vol) iron pyrophosphate. For mouse infections, *F. tularensis* was first grown on sMHA then transferred to Brain Heart Infusion agar (BHI; Becton Dickinson). Chocolate agar for mutant strain generation was prepared by mixing Mueller-Hinton broth powder with 1.6% (wt/vol) agar, 1% (wt/vol) tryptone, and 0.5% (wt/vol) sodium chloride, autoclaved, and further supplemented with 1% (wt/vol) hemoglobin and 1% (vol/vol) IsoVitaleX. For macrophage infections, *F. tularensis* was first grown on sMHA then transferred to modified chocolate agar: Mueller-Hinton broth powder was mixed with 1.6% (wt/vol) Bacto Agar, 1% hemoglobin (wt/vol), and 1% (vol/vol) IsoVitaleX. All growth curves were performed in sMHB: Mueller-Hinton broth powder was mixed with 182 µg ml^-1^ calcium chloride dihydrate, and 210 µg ml^-1^ magnesium chloride hexahydrate, 0.1% (wt/vol) glucose, 0.025% (wt/vol) iron pyrophosphate, and 2% (vol/vol) IsoVitaleX. All bacterial strains and plasmids are listed in Table S7. *E. coli* S17-1 and *E. coli* NEB10-β were grown in Luria Bertani (LB) broth or on LB agar at 37°C, supplemented as needed with antibiotics.

### Sequence alignments and bioinformatic predictions

Amino acid alignments of *F. tularensis* subsp. *holarctica* FTL_1678, *F. tularensis* subsp. *tularensis* FTT_0101, *E. coli* LdcA (BAA36050.1), *P. aeruginosa* LdcA (Q9HTZ1), *N. gonorrhoeae* LdcA (YP_208343.1), and *C. jejuni* Pgp2 (WP_002856863) were performed using Clustal Omega (https://www.ebi.ac.uk/Tools/msa/clustalo/) and MView (https://www.ebi.ac.uk/Tools/msa/mview/). Pairwise sequence alignments were performed and amino acid identities among Ldc homologs were calculated by EMBOSS Needle (https://www.ebi.ac.uk/Tools/psa/emboss_needle/). The Prokaryotic Genome Analysis Tool (PGAT) (http://tools.uwgenomics.org/pgat/), BlastP, and BlastX analyses (http://blast.ncbi.nlm.nih.gov) were used to identify *F. tularensis* homologues. Bacterial protein sub-localization was predicted by PSORTb version 3.0.2 (https://www.psort.org/psortb/). Protein signal sequence prediction was performed by LipoP version 1.0 (http://www.cbs.dtu.dk/services/LipoP/) and SignalP version 4.1 (http://www.cbs.dtu.dk/services/SignalP-4.1/).

### Expression and purification of recombinant FTL1678 and FTT0101

*F. tularensis* LVS and SchuS4 genomic DNA were extracted using phenol/chloroform/isoamyl alcohol (Fisher Bioreagents). *FTL1678* and *FTT0101*, without signal sequences (amino acid residues 1-29), were PCR-amplified from LVS and SchuS4 genomic DNA, respectively, using High Fidelity Platinum Taq Polymerase (Life Technologies), and primers 5’FTL1678_BamHI and 3’FTL1678_XhoI and 5’FTT0101_BamHI and 3’FTT0101_XhoI, respectively (Table S8). Amplicons and pPROEX HTb were double-digested with BamHI and XhoI, ligated using T4 DNA ligase, and transformed into NEB 10-β *E. coli*. Plasmids were purified using the Qiagen QIAprep Spin Miniprep kit and diagnostic PCR was performed to confirm presence and correct size of the insert. DNA sequencing was performed to confirm insert integrity and plasmid constructs were transformed into Rosetta DE3 *E. coli* (Millipore) for protein expression. Recombinant proteins were expressed and purified as previously described (Ren, 2014) with some modifications. Bacteria were grown in LB-amp to an OD_600_ of 0.4, protein expression was induced for 2 h by the addition of isopropyl β-D-thiogalactopyranoside (IPTG) to a final concentration of 100 µM, bacteria were pelleted by centrifugation, and frozen overnight at −80°C to aid in lysis. Cell pellets were suspended in 10 mM Tris, 500 mM NaCl, and 10 mM imidazole, pH 8.0, sonicated on ice for 10 min with 30 sec intervals, insoluble material was removed by centrifugation at 8,000 × *g*, and supernatants were collected for affinity purification over pre-equilibrated nickel-nitrilotriacetic acid (Ni-NTA) agarose (Qiagen) columns. Eluted recombinant proteins were concentrated in Amicon Ultra-4 centrifugal filter units with 30-kDa cutoff (Millipore), concentrations were determined using the DC BCA protein assay (BioRad), and purity was assessed by SDS-PAGE and Imperial protein staining (Thermo Scientific). An empty vector construct also was expressed and purified as a control in enzymatic assays.

### Site-directed mutagenesis of recombinant FTL1678

The QuikChange Lightning site-directed mutagenesis kit (Agilent) was used to generate single amino acid, double amino acid, and triple amino acid mutations in the FTL1678 putative active site residues S134, E239, and H308. Three separate QuikChange reactions were performed in a thermocycler using purified plasmid DNA from pFNLTP6-gro-*flt1678*-6xHis and three primer sets to individually mutate Ser134 to Ala (S134A), Glu239 to Ala (E239A), and His308 to Ala (H308A): a400g_g401c_5’ and a400g_g401c_3’(S134), a716c_5’ and a716c_3’ (E239), and c922g_a923c_5’ and c922g_a923c_3’(H308) (Table S8). Following amplification, products were digested with DpnI and immediately transformed into NEB 10-β *E. coli*. Transformants were selected on LB-kan overnight, plasmids were purified for individual clones, and DNA sequencing was performed to confirm individual mutations. The resulting plasmids were named pFNLTP6-gro-*ftl1678*-S134A-6xHis, pFNLTP6-gro-*ftl1678*-E239A-6xHis, and pFNLTP6-gro-*ftl1678*-H308A-6xHis (Table S7). To generate the catalytic triad point mutant complement strains, S134A, E239A, and H308A, pFNLTP6-gro-*ftl1678*-S134A-6xHis, pFNLTP6-gro-*ftl1678*-E239A-6xHis, and pFNLTP6-gro-*ftl1678*-H308A-6xHis were individually transformed Δ*FTL1678* by electroporation, transformants were selected on sMHA-kan10, and expression of S134A, E239A, and H308A were confirmed by immunoblot analysis. To generate double and triple active site mutants for recombinant protein expression, three separate QuikChange reactions were performed using purified plasmid DNA from pPROEX Htb-FTL1678 as described above, using the same three primer sets to mutate S134, E239, and H308. DNA sequencing was performed to confirm individual mutations. Plasmids were named pPROEX Htb-*ftl1678*-S134A, pPROEX Htb-*ftl1678*-E239A, and pPROEX Htb-*ftl1678*-H308A (Table S7). Three separate QuikChange reactions were performed as described above to generate double mutants. Purified plasmid DNA from pPROEX Htb-*ftl1678*-S134A was amplified with a716c_5’ and a716c_3’ to generate an S134A/E239A double mutant and separately with c922g_a923c_5’ and c922g_a923c_3’ to generate an S134A/H308A double mutant. Purified plasmid DNA from pPROEX Htb-*ftl1678*-E239A was amplified with c922g_a923c_5’ and c922g_a923c_3’ to generate an E239A/H308A double mutant. Transformants were selected on LB-kan overnight, plasmids were purified for individual clones, and DNA sequencing was performed to confirm double mutations. The resulting plasmids were named pPROEX Htb-*ftl1678*-S134A/E239A, pPROEX Htb-*ftl1678*-S134A/H308A, and pPROEX Htb-*ftl1678*-E239A/H308A. A final QuikChange reaction was performed to generate the triple active site mutant, S134A/E239A/H308A. Purified plasmid DNA from pPROEX Htb-*ftl1678*-S134A/E239A was amplified using the primer set c922g_a923c_5’ and c922g_a923c_3’, transformants were selected on LB-kan overnight, plasmids were purified for individual clones, DNA sequencing was performed to confirm the triple mutation and the plasmid was named pPROEX Htb-*ftl1678*-S134A/E239A/H308A (Table S7). All four plasmid constructs were transformed into Rosetta DE3 *E. coli* for protein expression. Recombinant proteins were expressed and purified as described above.

### Enzymatic assays for FTL1678/FTT0101 activity

FTL1678 and FTT0101 recombinant protein activities toward various PG-related compounds were tested in 50 µl reaction mixtures containing 50 mM Tris-HCl, pH 8.0, 0.1 mM substrate, and partially purified enzyme stock (10 µl in 1 M NaCl, 10 mM Tris, pH 8.0). Mixtures were incubated at 37°C and reactions were stopped by freezing. Depending on the substrate used, the amount of partially purified protein varied from 0.9 to 5 µg per assay and incubation time varied from 30 min to 4 h. Substrate and reaction products were separated by HPLC on an ODS-Hypersil 3 µm particle-size C18 column (250 × 4.6 mm; Thermo Scientific). Elutions were performed with 50 mM sodium phosphate buffer, pH 4.5, with or without application of a linear gradient of methanol (from 0 to 20% in 80 min), at a flow rate of 0.5 ml/min. Peaks were detected by measuring the absorbance at 207 nm or at 262 nm for UDP-containing nucleotide precursors. Identification of reaction products was based on their retention times, compared to authentic standards, as well as on their amino acid and amino sugar composition, determined with a Hitachi model L8800 analyzer (Sciencetec) after hydrolysis of samples in 6 M HCl for 16 h at 95°C. Enzyme activity was calculated by integration of peaks corresponding to the substrates and products. To ensure linearity, substrate consumption was < 20% in all cases. The amounts of D-Ala and D-Ala-D-Ala released during the reactions also were determined by injection of aliquots of these reaction mixtures in the Hitachi amino acid analyzer. FTL1678 double and triple active site mutants were tested using the same conditions, with the incubation time being prolonged up to 18h.

### Peptidoglycan precursors and muropeptides

UDP-MurNAc-pentapeptide precursors containing either *meso*-diaminopimelic acid (A_2_pm) or L-lysine were prepared by enzymatic synthesis using purified Mur ligases, and UDP-MurNAc-tetrapeptides were generated by treatment of the UDP-MurNAc-pentapeptide precursors with purified *E. coli* PBP5 DD-carboxypeptidase as previously described (Herve, 2007). MurNAc-peptides were obtained by mild acid hydrolysis of UDP-MurNAc-peptides (0.1 M HCl, 100°C, 15 min) and were not reduced and thus purified as a mixture of the two α and β anomers (Blanot, 1983). Free peptides were prepared by cleavage of MurNAc-peptides with *E. coli* AmiD *N*-acetylmuramoyl-L-alanine amidase (Pennartz, 2009). The *E. coli* peptidoglycan polymer was purified from a *lpp* mutant strain that does not express the Lpp lipoprotein (Leulier, 2003). GlcNAc-1,6-anhydro-MurNAc-L-Ala-γ-D-Glu-*meso*-A_2_pm-D-Ala (TCT) and its dimer (two cross-linked TCT monomers) were produced by digestion of peptidoglycan with *E. coli* SltY lytic transglycosylase and the non-anhydro forms of these monomer and dimer were generated by digestion of the polymer with mutanolysin (Stenbak, 2004). All these compounds were HPLC-purified (Table S1) and their composition was controlled by amino acid and amino sugar content analysis and/or by MALDI-TOF mass spectrometry.

### Generation of F. tularensis gene deletion mutants

*F. tularensis* isogenic deletion mutants were generated by homologous recombination as previously described (Wu, 2015). Briefly, 500-bp regions upstream and downstream from the gene of interest (*FTL1678* or *FTT0101*) were PCR-amplified from *F. tularensis* genomic DNA using the following primers: FTL1678_A and FTL1678_B; FTL1678_C and FTL1678_D; FTT0101_A and FTT0101_B; FTT0101_C and FTT0101_D (Table S8). A FLP recombination target (FRT)-flanked Pfn-kanamycin resistance cassette, FRT-Pfn-kan-FRT, was PCR amplified from pLG66a (Gallagher, 2008) and splicing overlap extension PCR (SOE PCR) was used to join the upstream (A-B) and downstream (C-D) regions with FRT-Pfn-kan-FRT, which replaced the gene of interest. The resulting insert and a suicide plasmid, pTP163 (Robertson, 2013), were digested with ApaI (New England Biolabs), and ligated using T4 DNA ligase (New England Biolabs). Gene deletion constructs were transformed into NEB10-β *E. coli* (New England Biolabs), sequence-verified, transformed into *E. coli* S17-1, and conjugation was performed with *F. tularensis* LVS on sMHA plates. Conjugants were recovered on chocolate agar supplemented with 200 µg ml^-1^ hygromycin and 100 µg ml^-1^ polymyxin B. Individual mutants were selected by sequential plating on sMHA supplemented with 10 µg ml^-1^ kanamycin (sMHA-kan10), sMHA-kan10 with 8% (wt/vol) sucrose, and final replica plating onto sMHA containing either 200 µg ml^-1^ hygromycin (sMHA-hyg200) or sMHA-kan10. 780 Hyg-sensitive and kan-resistant 781 colonies were sequence verified (referred to hereafter as either Δ*FTL1678* or Δ*FTT0101*).

### FTL1678 complementation in trans

Complementation *in trans* was performed as previously described, with some modifications (Wu, 2016). *FTL1678* was PCR-amplified from *F. tularensis* LVS using primers 5’FTL1678_NEBuilder and 3’FTL1678_NEBuilder (Table S8), pQE-60 (Qiagen) was double-digested with NcoI and BglII (New England Biolabs), and the NEBuilder HiFi DNA Assembly Cloning kit was used to ligate the *FTL1678* amplicon and digested pQE-60. The construct was transformed into NEB 10-β *E. coli* and transformants were selected on LB agar supplemented with 100 µg ml^-1^ ampicillin (LB-amp). Plasmids were purified from individual clones using the Qiagen QIAprep Spin Miniprep kit (Qiagen), diagnostic PCR was performed to confirm insert presence and correct size, and DNA sequencing was performed to verify insert integrity. The resulting construct, *FTL1678* with a C-terminal 6×histidine tag, was PCR-amplified using primers 5’FTL1678_pFNLTP6 and 3’FTL1678_pFNLTP6 (Table S8), the amplicon and pFNLTP6-gro-GFP (Maier, 2004) were double-digested with XhoI and BamHI (New England Biolabs), and ligated using T4 DNA ligase. The construct, pFNLTP6-gro-Δ*FTL1678*-6xHis, was transformed into NEB10-β *E. coli*, transformants were selected on LB plates supplemented with 50 µg ml^-1^ kanamycin (LB-kan), and DNA sequencing was performed to verify FTL1678-6xHis integrity. Next, the kan resistance gene was removed from Δ*FTL1678* by suspending the strain in 0.5 M sucrose (in 1 mM EDTA, pH 7.5), washing three times, and electroporating the shuttle plasmid pTP405 (Robertson, 2013), which encodes the Flp recombinase to remove FRT-Pfn-kan-FRT from the genome. Bacteria were grown overnight on sMHA-hyg200, hyg-resistant transformants were passaged three times on sMHA, then transformants were replica plated onto sMHA-hyg200 and sMHA-kan10 to confirm sensitivity to both antibiotics (kan-cured Δ*FTL1678*). pFNLTP6-gro-Δ*FTL1678*-6×His was transformed into kan-cured *1678* by electroporation, transformants were selected on sMHA-kan10, and FTL1678 expression was confirmed by immunoblot analysis (referred to hereafter as Δ*FTL1678 trans*-complement).

### C. jejuni pgp2 complementation in trans

Complementation of *C. jejuni pgp2* (CJJ81176_0915) into Δ*FTL1678* was performed as described above, with several modifications. The *pgp2* gene, with the FTL1678 signal sequence (amino acid residues 1-29) in place of the native Pgp2 signal sequence (amino acid residues 1-18), was synthesized and inserted in pQE-60 by GenScript USA. pQE-60-*pgp2* was transformed into NEB10-β *E. coli* and selection was performed on LB-amp. Pgp2-6×His was amplified from pQE-60 using primers 5’FTL1678_pFNLTP6 and 3’FTL1678_pFNLTP6 (Table S8), the amplicon was ligated into similarly digested pFNLTP6, pFNLTP6-gro-*pgp2*-6×His was transformed into NEB10-β *E. coli*, and transformants were selected on LB-kan. Plasmids were purified from kan-resistant transformants, sequence verified, then electroporated into kan-cured Δ*FTL1678*. Pgp2 expression was confirmed by immunoblot analysis.

### Mouse infections

All animal studies were approved by the University of Toledo Institutional Animal Care and Use Committee (IACUC). Mouse infections were performed as previously described (Huntley, 2008), with some modifications. Briefly, *F. tularensis* strains were grown on sMHA overnight, transferred to BHI agar for an additional 20-24 h, suspended in sterile PBS, and diluted to the desired concentration (20 to 10^9^ CFU/20 µl) based on previous OD_600_ measurements and bacterial enumeration studies. Groups of 4-8 female C3H/HeN mice (6-8 weeks old; Charles River Laboratories) were anesthetized with a ketamine-xylazine sedative and intranasally (i.n.) infected with 20 µl of prepared bacterial suspensions. Bacterial inocula were serially-diluted and plated in quadruplet on sMHA to confirm CFUs. For survival studies, mice were monitored daily, for signs of disease, with health status scores (scale of 1-5, with 1 indicating healthy and 5 indicating mouse found dead) being recorded for each mouse. Moribund mice were humanely euthanized to minimize suffering. To quantitate bacterial tissue burdens, groups of 4 mice were euthanized on days 2 and 5 post-infection, blood was collected by cardiac puncture and plated onto sMHA, lungs, livers, and spleens were aseptically harvested, homogenized, 25 µl of PBS/mg of tissue was added to each tissue, serially-diluted, and dilutions were plated onto sMHA. Following 72 h of incubation, the number of colonies per plate were counted and CFU/mg (tissues) or CFU/ml (blood) were calculated based on tissue weight and dilution factor. For immunization and challenge studies, groups of 4-10 mice were i.n. immunized with either 100-300 CFU LVS or 10^4^-10^9^ CFU FTL1678, boosted 3-4 weeks later with either 10^3^ CFU LVS or 10^9^ CFU FTL1678, transported to the ABSL3 facility 3-weeks later, and i.n. challenged with 20-120 CFU of *F. tularensis* SchuS4. Mice were monitored daily for signs of disease with health status scores being recorded for each mouse.

### Membrane integrity testing

Sensitivity of LVS, Δ*FTL1678*, FTL1678 *trans*-complement, and the Pgp2 *trans*-complement to various antibiotics, detergents, dyes, and cell wall stressors was determined by disk diffusion assays or in liquid cultures, as previously described (Wu, 2016), with some modifications. Bacterial strains were grown on either sMHA or sMHA-kan10 (Δ*FTL1678* and complemented strains), scraped and resuspended in sterile PBS, adjusted to an OD_600_ of 0.2 (approx. 9 × 10^7^ CFU/ml), diluted 1:1 in PBS, and 100 µl was plated onto sMHA plates using cotton tipped applicators (Puritan). Sterile paper disks (Whatman; 0.8 mm thick, 6.5 mm in diameter) were placed in the center of each plate and antibiotics, detergents, or dyes were added to the disks at the concentrations listed in Table 2. Antibiotics tested were: gentamicin (Gibco), tetracycline (Fisher Scientific), chloramphenicol (Acros Organics), ciprofloxacin (Oxoid), ampicillin (Fisher Scientific), vancomycin (Acros Organics), bacitracin (Oxoid), bacitracin (Oxoid), ciprofloxacin (Oxoid), and polymyxin B (MP Biomedicals). Detergents tested were: sodium dodecyl sulfate (SDS; anionic; Fisher Scientific), Triton X-100 (nonionic; Acros Organics), cetyltrimethyl ammonium bromide (CTAB; cationic; MP Biomedicals), 3-cholamidopropyl dimethylammonio 1-propanesulfonate (CHAPS; zwitterionic; Thermo Scientific). In addition, sensitivity to ethidium bromide (Thermo Scientific) and lysozyme (Thermo Fisher) also was tested. After 48 h, diameters of zones of inhibition around the disks were measured, with experiments performed in triplicate to confirm reproducibility. Zones of inhibition were averaged, standard deviations calculated, and all data rounded to the nearest whole number. For liquid cultures, bacteria were suspended in sMHB, adjusted to OD_600_ 0.4, and 5 ml of each bacterial suspensions was inoculated into 100 ml of either sMHB or sMHB with 5 mM hydrogen peroxide (H_2_O_2_), 5% sodium chloride (NaCl), or pH 5.5 (pH of sMHB is 6.5). Cultures were grown in triplicate at 37°C with rotation at 180 rpm for 24 h with OD_600_ readings recorded every 4 h.

### Electron microscopy

Electron microscopy was used to visualize differences in bacterial envelope structure and cell shape, as previously described (Wu, 2016), with some modifications. WT LVS, Δ*FTL1678 trans*-complement (FTL1678 compl), and the Pgp2 *trans*-complement (Pgp2 compl) were grown overnight in sMHB, approx. 1 × 10^9^ CFU of each bacterial strain was pelleted by centrifugation at 7000 × *g* at 4°C, washed three times in PBS, fixed in 3% (vol/vol) glutaraldehyde (Electron Microscopy Sciences [EMS]) for approx. 24 h, washed twice in sodium cacodylate buffer (pH 7.4; EMS) for 10 min, suspended in 1% (wt/vol) osmium tetroxide (EMS) in s-collidine buffer (pH 7.4; EMS) for 45 min at room temperature (r/t) to stain and fix the samples, washed two times with sodium cacodylate buffer for 10 min each, and tertiary fixation was performed using an aqueous saturated solution of uranyl acetate (pH 3.3; EMS) for 45 min at r/t. Samples were then dehydrated at room temperature using a series of ethanol washes: two washes with 30% ethanol for 10 min each; two washes with 50% ethanol for 10 min each; two washes with 95% ethanol for 10 min each; two washes with 100% ethanol for 10 min each; and two washes with 100% acetone for 10 min each. Samples were then infiltrated with 50% acetone and 50% embedding media (Hard Plus Resin 812, EMS) for 8 h to overnight at r/t. Samples were embedded in 100% embedding media (EMS) and allowed to polymerize for 8 h to overnight at 85°C, then sectioned at 85-90 nm, and visualized using a Tecnai G2 Spirit transmission electron microscope (FEI) at 80 kV and Radius 1.3 (Olympus) imaging software at the University of Toledo Electron Microscopy Facility. For outer membrane (OM) thickness measurements, individual bacterial cells were analyzed at 120,000×, multiple measurements (3 to 7 per bacteria) of OM thickness per bacterium were calculated by Radius 1.3 imaging software using default settings, and average OM thickness per bacterial cell were recorded (WT n=50 bacterial cells; Δ*FTL1678* n=50 bacterial cells; FTL1678 compl n=44 bacterial cells; Pgp2 compl n=33 bacterial cells). For bacterial cell width measurements, individual bacterium were analyzed at 120,000×, cell length and width were calculated using default settings, and the smallest measurement (width) per bacterium was recorded (Figure 3: WT n=175 bacterial cells, Δ*FTL1678* n=175 bacterial cells; Figure S7: WT n=119 bacterial cells; Δ*FTL1678* n=120 bacterial cells; FTL1678 compl n=119 bacterial cells; Pgp2 compl n=119 bacterial cells). Experiments were performed twice to confirm reproducibility, with two bacterial preparations fixed, stained, embedded, sectioned, and visualized per experiment.

### Spheroplasting and sucrose density gradient centrifugation

Spheroplasting, osmotic lysis, and sucrose density gradient centrifugation was performed as previously described (Huntley, 2007) to determine subcellular localization of FTL1678. Briefly, the histidine-tagged FTL1678 *trans*-complement was grown in sMHB to an OD_600_ of 0.3-0.4, pelleted at 7500 × *g* for 30 min at 10°C, supernatants were removed, pellets were resuspended in 0.75 M sucrose (in 5 mM Tris, pH 7.5) with gentle mixing, 10 mM EDTA (in 5 mM Tris, pH 7.8) was slowly added over 10 min, and the suspension was incubated for 30 min at r/t. After incubation, lysozyme was slowly added to a final concentration of 200 μ min at r/t, bacteria were osmotically lysed by dilution into 4.5× volumes of molecular-grade water (Corning) over 11 min with gentle mixing, and incubated for 30 min at r/t. Lysates were centrifuged at 7,500 × *g* for 30 min at 10°C to remove intact cells and cellular debris. Supernatants were collected and centrifuged at 182,500 × *g* for 2 h at 4°C in a F37L 8 × 100 Fiberlite Ultracentrifuge rotor. Following centrifugation, supernatants were removed, membrane pellets were gently resuspended in 6 ml of resuspension buffer (25% [wt/wt] sucrose, 5 mM Tris, 30 mM MgCl_2_, 1 tablet of Pierce Protease Inhibitor Mini Tablets, EDTA-Free [Thermo Scientific], 5 U Benzonase [Novagen]), suspensions were incubated with gentle mixing for 30 min at room temperature to degrade DNA, and a DC protein assay (Bio-Rad) was performed to determine total protein yield. Linear sucrose gradients were prepared by layering 1.8 ml each of sucrose solutions (wt/wt; prepared in 5 mM EDTA, pH 7.5) into 14-by 95-mm ultracentrifuge tubes (Beckman) in the following order: 55%, 50%, 45%, 40%, 35%, and 30%. Membrane suspensions were layered on top of each sucrose gradient, with less than 1.5 mg of protein per gradient. Sucrose gradients were centrifuged in an SW40 swinging bucket rotor (Beckman) at 256,000 × *g* for 17 h at 4°C. After centrifugation, 500-μ l fractions were collected from each gradient by puncturing the bottom of each tube and allowing fractions to drip into microcentrifuge tubes. The refractive index of each fraction was determined using a refractometer (Fisher Scientific) and correlated with a specific density in g ml^-1^ (Price, 1982) to identify outer membrane (OM; 1.17-1.20 g ml^-1^) and inner membrane (IM; 1.13-1.14 g ml^-1^) fractions. Sucrose gradient fractions were examined by immunoblotting as described below.

### Immunoblotting

Whole cell lysates of FTL1678 *trans*-complement were prepared by suspending bacteria (pelleted at 7000 × *g*) in molecular biology grade water, diluting with SDS-PAGE loading buffer, and boiling for 10 min. Whole cell lysates, OM fractions, IM fractions, and molecular mass standards (Precision Plus protein all blue prestained protein standards; BioRad Laboratories) were separated on a 12.5% polyacrylamide gel, transferred to nitrocellulose, and blots were incubated overnight in blot block (0.1% (vol/vol) Tween 20 and 2% (wt/vol) bovine serum albumin in PBS) at 4°C. Immunoblotting was performed using rat polyclonal antiserum specific for either *F. tularensis* OM protein FopA, *F. tularensis* IM protein SecY (Huntley, 2007) or the Penta-His HRP conjugate antibody (Qiagen).

### Infections of mouse bone marrow derived macrophages (mBMDMs) and J774A.1 cells

Macrophage culture (37°C with 5% CO_2_ unless otherwise indicated) and infections were performed as previously described (Wu, 2016), with some modifications. Bone marrow macrophages were harvested from female C3H/HeN mice. Mice were euthanized by CO_2_ asphyxiation and cervical dislocation. Femurs and tibias of both hind legs were aseptically-harvested, marrow was flushed from each bone with RPMI-1640 (Hyclone) containing 10% heat-inactivated fetal bovine serum ([HI-FBS], Atlanta Biologicals) and 30% supernatants from day 7 L929 cultures (ATCC). Bone marrow was disrupted by repeated passage through a 23-gauge needle and cultured for 4 days. Next, cell media was removed and replaced with RPMI containing 10% HI-FBS and 30% supernatant from day 14 L929 cultures, and cells were cultured for 2 days. Approx. 24 h before infection, media was removed, cells were harvested by scraping and centrifugation at 400 × *g* for 10 min at 10°C, cells were enumerated using a hemocytometer, and diluted to 1×10^5^ cells in RPMI containing 10% HI-FBS. J774A.1 cells (ATCC) were cultured in Dulbecco’s Modified Eagle Medium ([DMEM], Gibco) containing 10% HI-FBS. Approx. 24 h before infection, cells were harvested as described above, seeded into individual wells of 24-well plates (Corning) at a concentration of 1×10^5^ cells/well, and incubated overnight. mBMDMs and J774A.1 cells were infected with a multiplicity of infection (MOI) of 100 bacteria to 1 cell (100:1). Following infection, cells were centrifuged at 1,000 × *g* for 10 min at 4°C, incubated at 37°C with 5% CO_2_ for 1 h, washed 1× with RPMI (or DMEM), media containing 100 µg ml^-1^ gentamicin was added to kill extracellular bacteria, cells were incubated at 37°C with 5% CO_2_ for 1 h, washed 1× with RPMI (or DMEM), lysed with 1% saponin for 4 min, serially diluted in PBS, plated onto sMHA plates, and bacteria were enumerated (entry) after 48 h. Alternatively, after gentamicin treatment and washing, RPMI (or DMEM) containing 10% HI-FBS was added to cells and they were incubated for 6 or 24 h, lysed, serially-diluted, and plated to determine bacterial numbers.

### Statistics

GraphPad Prism6 was used in various statistical analyses, including: differences in antibiotic, detergent, dye, or lysozyme susceptibility were calculated by one-way ANOVA with multiple comparisons and the Holm-Sidak post-hoc test; differences in EM measurements were determined by unpaired t-tests; differences in median time-to-death and percent survival following *F. tularensis* infection of mice were calculated using the log-rank Mantel-Cox test; differences in pathology scores of *F. tularensis*-infected tissues were calculated by two-way ANOVA with multiple comparisons and a Tukey post-hoc test. Differences in lung, liver, spleen, and blood bacterial burdens from infected mice were calculated by one-way ANOVA with multiple comparisons using R software.

## Supporting information

Zellner Supplemental Data

## Acknowledgments

We would like to acknowledge Drs. Joe Dillard and Ryan Schaub (University of Wisconsin-Madison) for all their help and expertise with peptidoglycan isolation/purification. We would also like to thank Dr. Erin Gaynor for her generous contribution of *C. jejuni* Pgp2-pGEM plasmid construct. This work was supported by grant R01 AI093351 from the National Institute of Allergy and Infectious Disease of the National Institutes of Health (NIAID-NIH) to J.F.H.

## Data Availability Statement

Data that support the findings of this study are available in the supplementary material of this article.

## Supporting Information

**Table S1.** HPLC retention times of peptidoglycan (PG) compounds analyzed in this study

**Table S2.** Endopeptidase activity of FTL1678

**Table S3.** Specific activity of FTL1678, FTL1678 double mutants, and FTL1678 triple mutants to the TCT monomer

**Table S4.** Bioinformatic analyses of FTL1678 localization

**Table S5.** Sensitivity of WT LVS, ⊗Δ*FTL1678*, *FTL1678 trans*-complement, and Pgp2 *trans*-complement to antibiotics, detergents, and dyes

**Table S6.** Sensitivity of WT *F. tularensis* SchuS4 and ⊗Δ*FTT0101* to antibiotics, detergents, and dyes

**Table S7.** Bacterial strains and plasmids used in this study

**Table S8.** Primers used in this study

**Figure S1. FTL1678 contains a putative L,D-carboxypeptidase domain.** NCBI Conserved Domain search results for *F. tularensis* FTL1678.

**Figure S2. Activity of FTL1678 and controls on GlcNAc-anhydroMurNAc-tetrapeptide (TCT).** (A) Structure of TCT with the LdcA cleavage site indicated (blue arrow). (B) HPLC analysis of reaction mixtures obtained following incubation of TCT in the presence of either FTL1678, vector control extract, or buffer alone. The enzymatic assay and HPLC conditions used are outlined in Materials and Methods.

**Figure S3. Activity of FTL1678 and controls on the free tetrapeptide, L-Ala-γ-D-Glu-*meso*-A_2_pm-D-Ala.** HPLC analysis of reaction mixtures obtained following incubation of L-Ala-γ-D-Glu-*meso*-A_2_pm-D-Ala (Tetra) in the presence of buffer alone, extract from the vector control, or purified FTL1678. A peak of L-Ala-γ-D-Glu-*meso*-A_2_pm product (Tri) was only detected in the presence of FTL1678. FTL1678 enzyme activity was not affected by the presence of either 2.5 mM MgCl_2_ or 5 mM EDTA added to the reaction mixtures. The enzymatic assay and HPLC conditions used are outlined in Materials and Methods.

**Figure S4. *F. tularensis* TolB is OM-localized.** Spheroplasting, osmotic lysis, and sucrose density gradient centrifugation were performed to separate inner membranes (IM) and outer membranes (OM) from *F*. *tularensis ΔFTL1678 trans*-complemented with a 6×histidine-tagged FTL1678. Whole-cell lysates (WCL), OM fractions, and IM fractions were separated by SDS-PAGE, transferred to nitrocellulose, and immunoblotting was performed using antisera specific for the periplasmic protein TolB (αTolB).

**Figure S5. Δ*FTL1678* does not have a growth defect.** WT and Δ*FTL1678* were grown in sMHB for 24 h at 37°C. Samples were taken every 4 h for (A) OD_600_ measurements and (B) CFU enumeration following serial-dilution and plating.

**Figure S6. Δ*FTL1678* has septation defects.** Transmission electron micrograph images of Δ*FTL1678* bacteria showing aberrant septal formation and reduced ability to separate cells. Images taken at: (A) 49,000×, scale bar represents 200 nm; and (B) 6,800×, scale bar represents 2 µm. In (A), red arrows point to formed septa that have not separated in Δ*FTL1678* bacteria and white arrows point to new septa that are forming in Δ*FTL1678* bacteria. Experiments were performed twice to confirm reproducibility, with two bacterial preparations fixed, stained, embedded, sectioned, and visualized per experiment. Representative images shown.

**Figure S7. *FTL1678 trans*-complement and *C. jejuni* Pgp2 *trans*-complement restore *F. tularensis* phenotype.** Transmission electron micrograph images of: (A) A) Δ*FTL1678 trans*-complemented with *FTL1678 trans*-complemented with Δ*FTL1678* [FTL1678 compl] or (B) *FTL1678 trans*-complemented with *C. jejuni pgp2* [Pgp2 compl]. Bacteria were grown in sMHB to OD_600_ of 0.4. Scale bars represent 100 nm. (C) Outer membrane thickness [WT n=50; Δ*FTL1678* n=50; FTL1678 compl n=44; Pgp2 compl n=33] and (D) cell width [WT n=119; Δ*FTL1678* n=120; FTL1678 compl n=119; Pgp2 compl n=119] of FTL1678 compl and Pgp2 compl were compared to WT LVS and *FTL1678.* **** indicates *P*<0.0001.

**Figure S8. *FTL1678 trans*-complement and *C. jejuni* Pgp2 *trans*-complement exhibit similar phenotypes to stressors as WT *F. tularensis*.** WT LVS [WT], *ΔFTL1678, ΔFTL1678 trans*-complemented with *FTL1678* [FTL1678 compl], or *ΔFTL1678 trans*-complemented with *C. jejuni pgp2* [Pgp2 compl] were grown in either: (A) sMHB; (B) sMHB with 5 mM H_2_O_2_; (C) sMHB with 5% NaCl; or (D) sMHB at pH 5.5. Cultures were incubated for 24 h and OD_600_ measurements were recorded every 4 h.

**Figure S9. FTT0101 is not required for *F. tularensis* Type A strain SchuS4 virulence.** C3H/HeN mice were intranasally infected with either 80 CFU SchuS4 (n=3 mice) or 12 CFU Δ*FTT0101* (n=5 mice).

**Figure S10. Individual amino acids of the LdcA catalytic triad are not essential for *F. tularensis* virulence.** Groups of 5 C3H/HeN mice were intranasally-infected with 10^5^ CFU of either *F. tularensis* WT LVS, Δ*FTL1678*, *ΔFTL1678 trans*-complemented with FTL1678 (FTL1678 compl), *ΔFTL1678 trans*-complemented with S134A (S134A), *ΔFTL1678 trans*-complemented with E239A (E239A), or *ΔFTL1678 trans*-complemented with H308A (H308A). Animal health was monitored daily through day 21 post-infection. ** *P*<0.01.

