## Supplementary material for "A *Francisella tularensis* L,D-carboxypeptidase plays important roles in cell morphology, envelope integrity, and virulence": Zellner Supplemental Data

**Summary:**
*Francisella tularensis* is a Gram-negative, intracellular bacterium that causes the zoonotic disease tularemia. Intracellular pathogens, including *F. tularensis*, have evolved mechanisms to survive in the harsh environment of macrophages and neutrophils, where they are exposed to cell envelope-damaging molecules. The bacterial cell wall, primarily composed of peptidoglycan (PG), maintains cell morphology, structure, and membrane integrity. Intracellular Gram-negative bacteria protect themselves from macrophage and neutrophil killing by recycling and repairing damaged PG – a process that involves over 50 different PG synthesis and recycling enzymes. Here, we identified a PG recycling enzyme, L,D-carboxypeptidase A (LdcA), of *F. tularensis* that is responsible for converting PG tetrapeptide stems to tripeptide stems. Unlike *E. coli* LdcA and most other orthologs, *F. tularensis* LdcA does not localize to the cytoplasm and also exhibits L,D-endopeptidase activity, converting PG pentapeptide stems to tripeptide stems. Loss of *F. tularensis* LdcA led to altered cell morphology and membrane integrity, as well as attenuation in a mouse pulmonary infection model and in primary and immortalized macrophages. Finally, an *F. tularensis ldcA* mutant protected mice against virulent Type A *F. tularensis* SchuS4 pulmonary challenge.

**Table S1.** HPLC retention times of peptidoglycan (PG) compounds analyzed in this study

| Substrates [a] | Retention time (min) [b] | Products [a] | Retention time (min) [b] |
|---|---|---|---|
| GlcNAc-anhydroMurNAc-tetrapeptide (TCT) | 52 | - GlcNAc-anhydroMurNAc-tripeptide<br>- D-Ala | 44<br>/ |
| GlcNAc-anhydroMurNAc-tetrapeptide (TCT) | 49 | - GlcNAc-anhydroMurNAc-tripeptide<br>- Gly | 44<br>/ |
| GlcNAc-MurNAc-tetrapeptide | 28/36 (α/β) [(c)] | - GlcNAc-MurNAc-tripeptide<br>- D-Ala | 21/29 (α/β)<br>/ |
| TCT dimer Tetra-Tetra ; 4-3 cross-link | 82 | - GlcNAc-anhydroMurNAc-tripeptide<br>- GlcNAc-anhydroMurNAc-tetrapeptide*<br>- D-Ala | 44<br>48<br>/ |
| TCT dimer Tri-Tri ; 3-3 cross-link | 73 | - GlcNAc-anhydroMurNAc-tripeptide (× 2) | 44<br>/ |
| TCT dimer Tri-Tetra ; 3-3 cross-link | 75 | - GlcNAc-anhydroMurNAc-tripeptide (× 2)<br>- D-Ala | 44<br>/ |
| TCT dimer Tri-Tetra($Gly_4$) ; 3-3 cross-link | 71 | - GlcNAc-anhydroMurNAc-tripeptide (× 2)<br>- Gly | 44<br>/ |
| MurNAc-tetrapeptide | 20/27 (α/β) | - MurNAc-tripeptide<br>- D-Ala | 15/21 (α/β)<br>/ |
| UDP-MurNAc-tetrapeptide | 21 | - UDP-MurNAc-tripeptide<br>- D-Ala | 15<br>/ |
| Tetrapeptide | 10.5 | - Tripeptide<br>- D-Ala | 9.1<br>/ |
| MurNAc-pentapeptide | 26/34 (α/β) | - MurNAc-tripeptide<br>- D-Ala-D-Ala | 15/21 (α/β)<br>10 |
| UDP-MurNAc-pentapeptide | 24.5 | - UDP-MurNAc-tripeptide<br>- D-Ala-D-Ala | 15<br>10 |
| MurNAc-tetrapeptide($Lys_3$) | 29/37 (α/β) | - MurNAc-tripeptide($Lys_3$) | 22/30 (α/β) |

| | | | |
|---|---|---|---|
| | | - D-Ala | / |
| Tetrapeptide($Lys_3$) | 15 | - Tripeptide($Lys_3$)<br>- D-Ala | 11<br>/ |
| UDP-MurNAc-pentapeptide($Lys_3$) | 31 | - UDP-MurNAc-tripeptide($Lys_3$)<br>- D-Ala-D-Ala | 22<br>10 |
| GlcNAc-MurNAc-tetrapeptide($A_2pm_{NH2}$) | 33/41 (α/β) | - GlcNAc-MurNAc-tripeptide($A_2pm_{NH2}$)<br>- D-Ala | 25/34 (α/β)<br>/ |
| GlcNAc-MurNAc-tetrapeptide($Glu_{NH2}$) | 31/39 (α/β) | - GlcNAc-MurNAc-tripeptide($Glu_{NH2}$)<br>- D-Ala | ND<br>/ |
| GlcNAc-MurNAc-tetrapeptide($Glu_{NH2}$/$A_2pm_{NH2}$) | 35/44 (α/β) | - GlcNAc-MurNAc-tripeptide($Glu_{NH2}$/$A_2pm_{NH2}$)<br>- D-Ala | ND<br>/ |

[a] Tetrapeptide or Tetra indicates the L-Ala-γ-D-Glu-*meso*-$A_2$pm-D-Ala peptide structure. ($Lys_3$) and ($Gly_4$) indicate that the $A_2$pm and D-Ala residues at positions 3 and 4 are replaced by lysine and glycine residues in these peptides, respectively. Tetrapeptide* indicates that the D-Ala residue is bound by its carboxyl group to the ε-amino group of $A_2$pm (and not by its amino group to the α-carboxyl group of $A_2$pm). ($Glu_{NH2}$) and ($A_2pm_{NH2}$) indicate that the α and ε carboxyl groups of these residues, respectively, are amidated.

[b] HPLC conditions used: column of ODS-Hypersil 3 µ (250 × 4.6 mm); elution with 50 mM sodium phosphate, pH 4.5, with a linear gradient of MeOH from 0 to 20% being applied between 0 and 80 min, at a flow rate of 0.5 ml/min; detection at 262 nm (nucleotide precursors) or 207 nm (other compounds). Enzyme activity is determined by integration of the peaks of substrates and products. The amounts of D-Ala and D-Ala-D-Ala released during the reactions were determined by injection of aliquots of these reaction mixtures on the Hitachi amino acid analyzer.

[c] Two peaks corresponding to the α and β anomers are observed for compounds containing a reducing MurNAc group.

[d] ND, not detected.

**Table S2.** Endopeptidase activity of FTL1678

| Substrate | Peptide length | Cross-linkage | Cleavage sites [a] | Products [b] | Specific activity (nmol/min/mg of protein)[c] |
|---|---|---|---|---|---|
| TCT Monomer | Tetra ($Ala_4$ ) | - | 1 (C) | TCT-tri + D-Ala | 20.1 ± 1.53 |
| TCT Monomer | Tetra ($Gly_4$ ) | - | 1 (C) | TCT-tri + Gly | 18.0 ± 1.84 |
| TCT Dimer | Tetra-Tetra | 4-3 (D,D) | 2 (E,C) | TCT-tri + TCT-tetra* + D-Ala | 6.6 ± 0.49 |
| TCT Dimer | Tri-Tri | 3-3 (L,D) | 1 (E) | TCT-tri + TCT-tri | 0.28 ± 0.051 |
| TCT Dimer | Tri-Tetra ($Ala_4$ ) | 3-3 (L,D) | 2 (E,C) | TCT-tri + TCT-tri +D-Ala | 0.16 ± 0.029 |
| TCT Dimer | Tri-Tetra ($Gly_4$ ) | 3-3 (L,D) | 2 (E,C) | TCT-tri + TCT-tri + Gly | 0.21 ± 0.042 |

[a] C indicates L,D-carboxypeptidase activity, E indicates L,D-endopeptidase activity.

[b] In TCT-tetra*, the D-Ala residue is bound by its carboxyl group to the ε-amino group of $A_2pm$.

[c] When two cleavage sites are present in the substrate, the indicated activity values correspond to the first and limiting reaction (endopeptidase) catalyzed by the enzyme. Values represent the mean ± SD of triplicate experiments.

**Table S3.** Specific activity of FTL1678, FTL1678 double mutants, and FTL1678 triple mutants to the TCT monomer

| **Enzyme** | **Specific activity (nmol/min/mg of protein) to TCT monomer**[a] |
|---|---|
| WT FTL1678 | 21.5 ± 1.34 |
| S134A/E239A | Not detected [b] |
| S134A/H308A | Not detected [b] |
| E239A/H308A | Not detected [b] |
| S134A/E239A/H308A | Not detected [b] |

[a] Standard enzyme assay conditions are described in the Materials and Methods. TCT monomer is GlcNAc-anhydroMurNAc-L-Ala-γ-D-Glu-*meso*-$A_2$pm-D-Ala.

[b] Not detected indicates that no release of alanine was detected.

**Table S4.** Bioinformatic analyses of FTL1678 localization

| Protein | PSORTb localization[a] | BOMP Score[b] | LipoP Score[c] | SignalP Score[d] |
|---|---|---|---|---|
| FTL1678 | Unk | 0 | SpI score = 9.29708<br>SpII score = 2.64664 | D score = 0.820,<br>position 1-27 |

[a] PSORTb version 3.0.2 bacterial protein subcellular localization prediction program (http://www.psort.org/psortb/). Outer membrane (OM), cytoplasmic (CYT), cytoplasmic membrane (CM), periplasmic (P), extracellular (EXT) or unknown (Unk) localization. Scores > 7.5 considered significant.

[b] BOMP beta-barrel integral outer membrane protein prediction program (http://services.cbu.uib.no/tools/bomp). Scores > 2 considered significant.

[c] LipoP 1.0 lipoprotein signal peptide prediction program (http://www.cbs.dtu.dk/services/LipoP/). SpI indicates signal peptidase I, SpII indicates signal peptidase II (lipoprotein signal peptide). Scores > 4.0 considered significant.

[d] SignalP 4.1 signal peptide prediction program (http://www.cbs.dtu.dk/services/SignalP-4.1/). D score >0.450 is considered significant. Position indicates signal peptide residues.

**Table S5.** Sensitivity of WT LVS, Δ*FTL1678*, FTL1678 *trans*-complement, and Pgp2 *trans*-complement to antibiotics, detergents, and dyes

| | | **Average zone of inhibition, mm (mean ± SD)** | | | |
|---|---|---|---|---|---|
| **Compound** | **Concentration (µg/disk)** | **WT LVS** | **Δ*FTL1678*** [a, b] | **FTL1678 *trans*-complement** | **Pgp2 *trans*-complement** |
| Gentamicin | 4 | 3 ± 0 | 1 ± 0 (R) | 3 ± 0 | 2 ± 0 |
| Tetracycline | 5 | 2 ± 0 | 1 ± 0 (R) | 2 ± 0 | 2 ± 0 |
| Chloramphenicol | 5 | 3 ± 0 | 1 ± 0 (R) | 3 ± 0 | 3 ± 0 |
| Ciprofloxacin | 5 | 5 ± 0 | 2 ± 0 (R) | 5 ± 0 | 5 ± 0 |
| Ampicillin | 200 | 3 ± 0 | 4 ± 0 (S) | 3 ± 0 | 3 ± 0 |
| Vancomycin | 20 | 1 ± 0 | 3 ± 0 (S) | 2 ± 0 | 2 ± 0 |
| Bacitracin | 182 | 1 ± 0 | 1 ± 0 | 2 ± 0 | 2 ± 0 |
| Polymyxin B | 100 | 1 ± 0 | 1 ± 0 | 1 ± 0 | 1 ± 0 |
| Lysozyme | 1000 | 1 ± 0 | 2 ± 0 (S) | 1 ± 0 | 1 ± 0 |
| Ethidium Bromide | 5 | 3 ± 0 | 1 ± 0 (R) | 3 ± 0 | 3 ± 0 |
| Triton-X 100 | 750 | 3 ± 0 | 3 ± 0 | 4 ± 0 | 4 ± 0 |
| SDS | 1000 | 1 ± 0 | 2 ± 0 (S) | 1 ± 0 | 1 ± 0 |
| CTAB | 50 | 1 ± 0 | 1 ± 0 | 1 ± 0 | 1 ± 0 |
| CHAPS | 50 | 1 ± 0 | 1 ± 0 | 1 ± 0 | 1 ± 0 |

[a] R indicates a strain is significantly more resistant than WT LVS by one-way ANOVA ($P$<0.05)

[b] S indicates a strain is significantly more sensitive than WT LVS by one-way ANOVA ($P$<0.05)

**Table S6.** Sensitivity of WT *F. tularensis* SchuS4 and Δ*FTT0101* to antibiotics, detergents, and dyes

| | | **Average zone of inhibition, mm (mean ± SD)** | |
|---|---|---|---|
| **Compound** | **Concentration (µg/disk)** | **WT SchuS4** | **Δ*FTT0101*** |
| Gentamicin | 4 | 2 ± 0 | 2 ± 0 |
| Tetracycline | 5 | 2 ± 0 | 2 ± 0 |
| Chloramphenicol | 5 | 2 ± 0 | 2 ± 0 |
| Ampicillin | 200 | 1 ± 0 | 1 ± 0 |
| Vancomycin | 20 | 1 ± 0 | 1 ± 0 |
| Bacitracin | 182 | 1 ± 0 | 1 ± 0 |
| Polymyxin B | 100 | 1 ± 0 | 1 ± 0 |
| Lysozyme | 1000 | 1 ± 0 | 1 ± 0 |
| Ethidium Bromide | 5 | 1 ± 0 | 2 ± 0 |
| Triton-X 100 | 750 | 3 ± 0 | 3 ± 0 |
| SDS | 1000 | 1 ± 0 | 1 ± 0 |
| CTAB | 50 | 1 ± 0 | 1 ± 0 |
| CHAPS | 50 | 1 ± 0 | 1 ± 0 |

**Table S7.** Bacterial strains and plasmids used in this study

| Strain or plasmid | Relevant genotype or strain information | Source |
|---|---|---|
| S17-1 | *E. coli* for conjugation | Simon *et al.*, 1983 |
| NEB10-β | *E. coli* chemically competent cells for transformation | New England Biolabs |
| LVS | *F. tularensis* subsp. *holarctica*, Type B, Live Vaccine Strain | BEI Resources |
| SchuS4 | *F. tularensis* subsp. *tularensis*, Type A | BEI Resources |
| Δ*FTL1678* | *FTL1678* isogenic deletion in LVS | This study |
| FTL1678 *trans* comp. | 6×His-tagged FTL1678 complemented in-*trans* in Δ*FTL1678* | This study |
| Pgp2 *trans* comp. | 6×His-tagged Pgp2 from *Campylobacter jejuni* complemented in-*trans* in Δ*FTL1678* | This study |
| Δ*FTT0101* | *FTT0101* isogenic deletion in SchuS4 | This study |
| pPROEX-Htb | pPROEX-Htb expression plasmid with N-term 6×His tag | Thermo Fisher |
| pQE-60 | pQE-60 expression plasmid with C-term 6×His tag | Qiagen |
| pLG66a | Source of FRT-Pfn-*kan*-FRT*,* kanamycin resistance, gift from C. Manoil, University of Washington | Gallagher *et al.*, 2008 |
| pTP163 | Suicide plasmid with *oriT* from pUC18T-mini-Tn*7*T and hygromycin resistance, gift from G. Robertson and M. Norgard, University of Texas Southwestern Medical Center | Robertson *et al.*, 2013 |
| pFNLTP6-gro-GFP | Green fluorescence protein expression plasmid for *Francisella* with a *groEL* promoter, gift from T. Zahrt, Medical College of Wisconsin | Maier *et al.*, 2004 |
| pQE-60-*FTL1678* | Full-length *FTL1678* | This study |
| pFNLTP6-gro-*FTL1678*-6xHis | Full-length *FTL1678* with a C-terminal 6×His tag | This study |
| pFNLTP6-gro-*pgp2*-6×His | Pgp2 from *C. jejuni* with the *FTL1678* signal sequence (residues 1-29) in place of the native signal sequence | This study |
| pPROEX Htb-*FTL1678* | *FTL1678* with signal sequence (amino acid residues 1-29) removed , N-terminal 6×His tag | This study |
| pPROEX Htb-*FTT0101* | FTT0101 with signal sequence (amino acid residues 1-29) removed , N-terminal 6× His tag | This study |
| pFNLTP6-gro-*ftl1678*-S134A-6xHis | Full-length FTL1678 with S134A mutation, C-terminal 6X Histidine tag | This study |
| pFNLTP6-gro-*ftl1678*-E239A-6xHis | Full-length FTL1678 with E239A mutation, C-terminal 6X Histidine tag | This study |
| pFNLTP6-gro-*ftl1678*-H308A-6xHis | Full-length FTL1678 with H308A mutation, C-terminal 6X Histidine tag | This study |
| pPROEX Htb-*ftl1678*-S134A | FTL1678 (signal seq. removed) with S134A mutation, N-terminal 6X Histidine tag | This study |

| | | |
|---|---|---|
| pPROEX Htb-*ftl1678*-E239A | FTL1678 (signal seq. removed) with E239A mutation, N-terminal 6X Histidine tag | This study |
| pPROEX Htb-*ftl1678*-H308A | FTL1678 (signal seq. removed) with H308A mutation, N-terminal 6X Histidine tag | This study |
| pPROEX Htb-*ftl1678*-S134A/E239A | FTL1678 (signal seq. removed) with S134A/E239A double mutation, N-terminal 6X Histidine tag | This study |
| pPROEX Htb-*ftl1678*-S134A/H308A | FTL1678 (signal seq. removed) with S134A/H308A double mutation, N-terminal 6X Histidine tag | This study |
| pPROEX Htb-*ftl1678*-E239A/H308A | FTL1678 (signal seq. removed) with E239A/H308A double mutation, N-terminal 6X Histidine tag | This study |
| pPROEX Htb-*ftl1678*-S134A/E239A/ H308A | FTL1678 (signal seq. removed) with S134A/E239A/H308A triple mutation, N-terminal 6X Histidine tag | This study |

**Table S8.** Primers used in this study

| Primer name | Sequence |
|---|---|
| FTL1678_A | 5'-GGTAAGGGGCCCCAGGGCAAAATACTCGAAAACAG-3' |
| FTL1678_B | 5'-AACTTCCGCCGGCGTAGTAGTTTATCTTAATTATAACATTAATATGGACATTAGATATT-3' |
| FTL1678_C | 5'-TTAGATGTGTATAAGCAGCCCGATTTAATCAAAGCGGAAATTTAGCC-3' |
| FTL1678_D | 5'-GCTACAGGGCCCCCGAACTGAGGTTATTTTAGCAGATAGT-3' |
| Kan 5’ | 5'-TACTACGCCGGCGGAAGTTCCTATAC-3' |
| Kan 3’ | 5'-GGCTGCTTATACACATCTAAGAAGTTCCTATTCC-3' |
| 5’FTL1678_NEBuilder | 5’-GAATTCATTAAAGAGGAGAAATTAACCATGGCATGTTGTTAAAAAAT-3’ |
| 3’FTL1678_NEBuilder | 5’-CTTAGTGATGGTGATGGTGATGAGATCTTTTTTGTTTTAAAGTG-3’ |
| 5’FTL1678_pFNLTP6 | 5’-GCGCCTCGAGATGTTGTTAAAAAATAATTATTTAGTAAGTATTATATTAGTAG-3’ |
| 3’FTL1678_pFNLTP6 | 5’-GCGCGGATCCTTAGTGATGGTGATGGTGATG-3’ |
| 5’FTL1678_BamHI | 5’-GCGCGCGGATCCGATTACAATAAAGTCGCTTTAATTAATG-3’ |
| 3’FTL1678_XhoI | 5’-GCGCGCCTCGAGTCATTATTTTTGTTTTAAAGTGCAAAATATAGTTT-3’ |
| FTT0101_A | 5’-GGTAAGGGGCCCACTGATGGCTGGAAAGC-3’ |
| FTT0101_B | 5’-AACTTCCGCCGGCGTAGTACAACATAAAGAATTTATTTATTAATAATGTTTATCTTAATT-3’ |
| FTT0101_C | 5’-TTAGATGTGTATAAGCAGCCCGATTTAATCAAAGCGGAAATTTAGCTTC-3’ |
| FTT0101_D | 5’-GCTACAGGGCCCCACACATTATCGAACAAATTTTATATCTTGTAACAAATG-3’ |
| 5’FTT0101_BamHI | 5’-GCGCGGATCCGATTACAATAAAGTCGCTTTAATTAATG-3’ |
| 3’FTT0101_XhoI | 5’-GCGCCTCGAGTCATTATTTTTGTTTTAAAGTGCAAAATATAG-3’ |
| a400g_g401c_5’ | 5'-AAAATGTATAGCTGTAACATCAGCAAATCCAACTAATATTTTAGGTTTAGCTTTTTTAAGCTCA-3' |
| a400g_g401c_3’ | 5'-TGAGCTTAAAAAAGCTAAACCTAAAATATTAGTTGGATTTGCTGATGTTACAGCTATACATTT-3' |
| a716c_5’ | 5'-CTGTCTAAATGAAATGCCAGTATCTGCAAGAAATAGAATTTTTGTTGAGAT-3' |
| a716c_3’ | 5'-ATCTCAACAAAAATTCTATTTCTTGCAGATACTGGCATTTCATTTAGACAG-3' |
| c922g_a923c_5’ | 5'-AAAGGTTTATTGTATTGTCCAGCACCTATAAAAGGAAAATAATATACAGGTCTATTAAAAGTTT-3' |
| c922g_a923c_3’ | 5'-AAACTTTTAATAGACCTGTATATTATTTTCCTTTTATAGGTGCTGGACAATACAATAAACCTTT-3' |

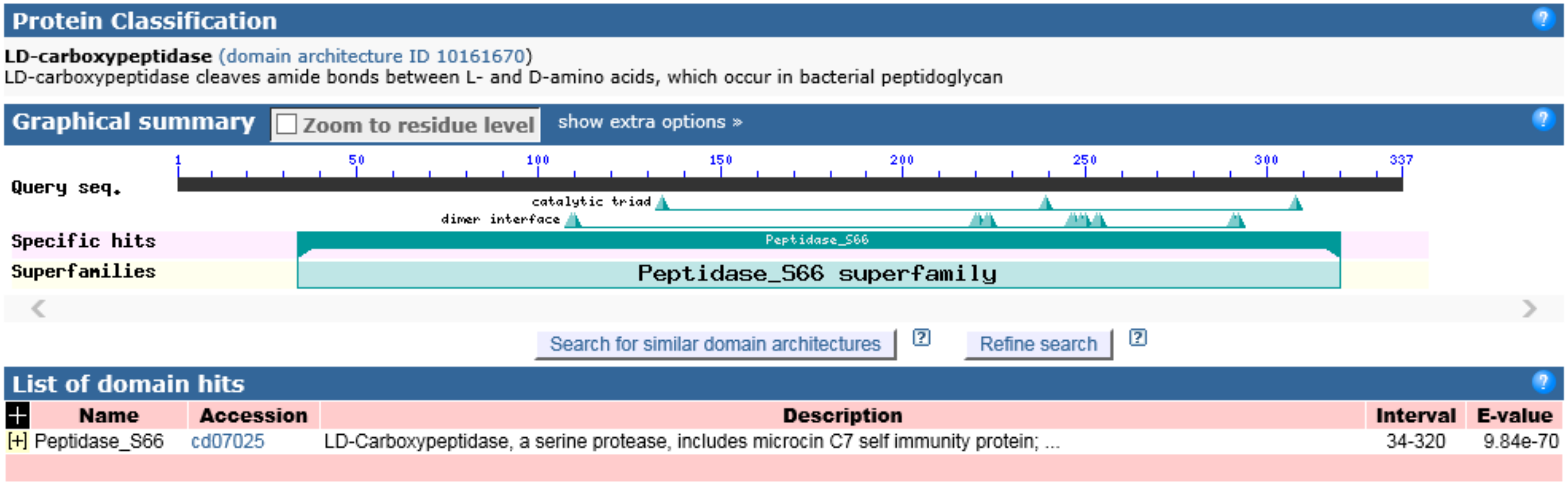


**Figure S1. FTL1678 contains a putative L,D-carboxypeptidase domain.** NCBI Conserved Domain search results for *F. tularensis* FTL1678.

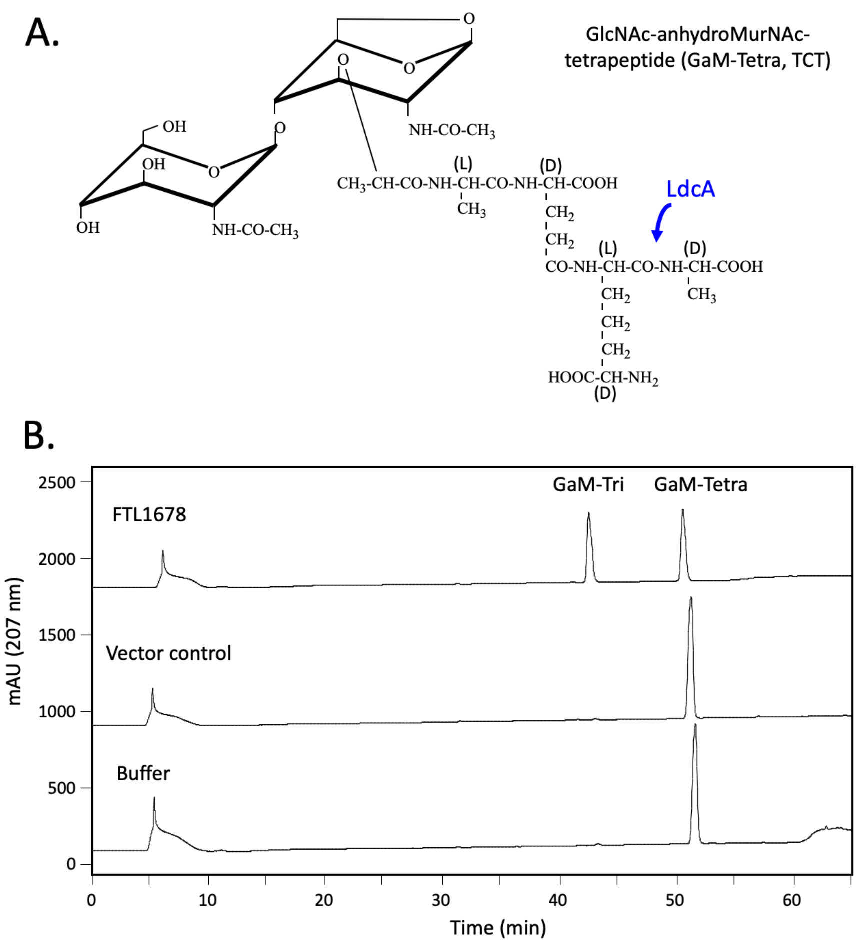


**Figure S2. Activity of FTL1678 and controls on GlcNAc-anhydroMurNAc-tetrapeptide (TCT).** (A) Structure of TCT with the LdcA cleavage site indicated (blue arrow). (B) HPLC analysis of reaction mixtures obtained following incubation of TCT in the presence of either FTL1678, vector control extract, or buffer alone. The enzymatic assay and HPLC conditions used are outlined in Materials and Methods.

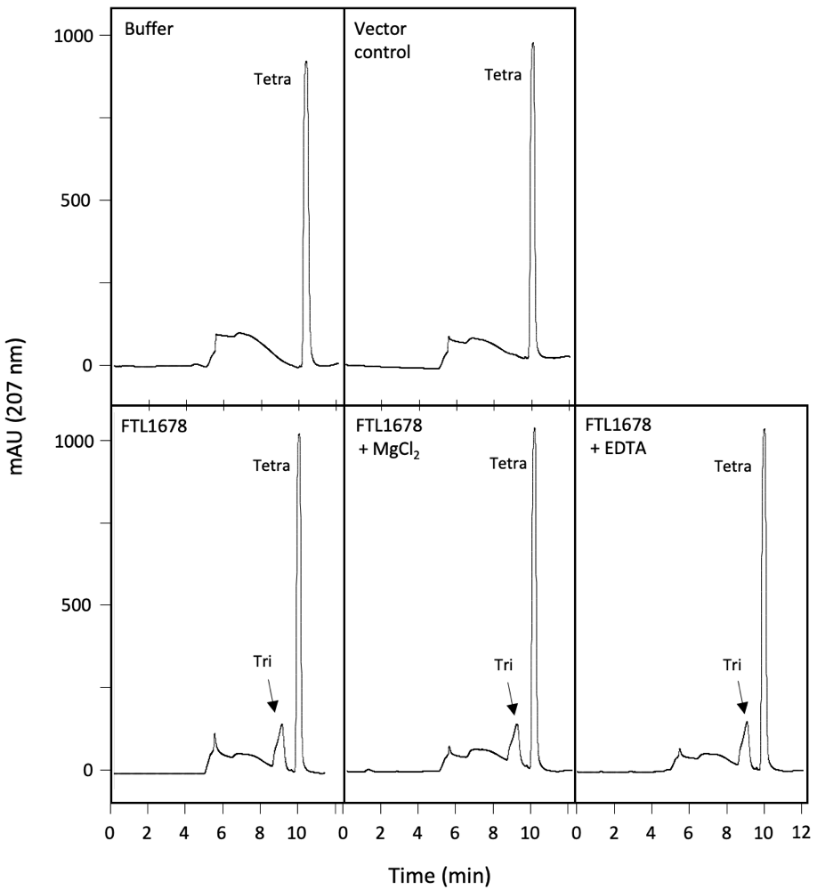


**Figure S3. Activity of FTL1678 and controls on the free tetrapeptide, L-Ala-γ-D-Glu-*meso*-$A_2$pm-D-Ala.** HPLC analysis of reaction mixtures obtained following incubation of L-Ala-γ-D-Glu-*meso*-$A_2$pm-D-Ala (Tetra) in the presence of buffer alone, extract from the vector control, or purified FTL1678. A peak of L-Ala-γ-D-Glu-*meso*-$A_2$pm product (Tri) was only detected in the presence of FTL1678. FTL1678 enzyme activity was not affected by the presence of either 2.5 mM $MgCl_2$ or 5 mM EDTA added to the reaction mixtures. The enzymatic assay and HPLC conditions used are outlined in Materials and Methods.

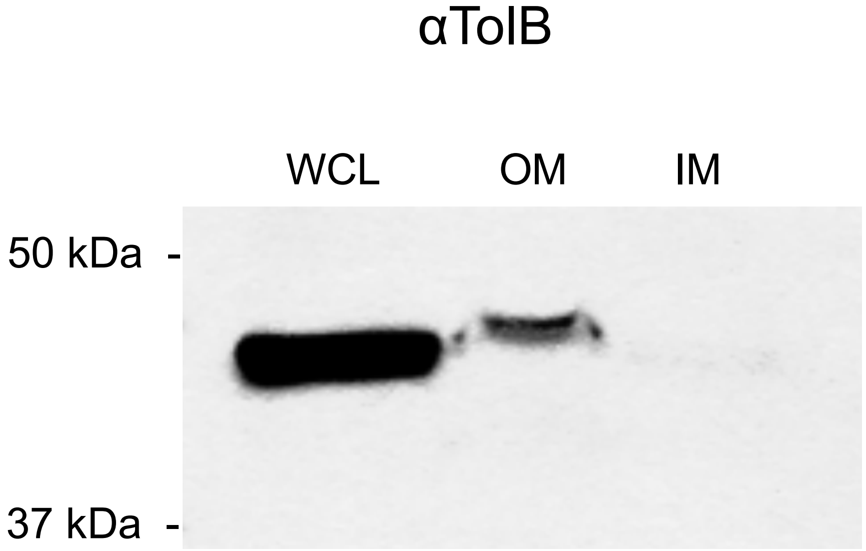


**Figure S4.** ***F. tularensis* TolB is OM-localized.** Spheroplasting, osmotic lysis, and sucrose density gradient centrifugation were performed to separate inner membranes (IM) and outer membranes (OM) from *F. tularensis* Δ*FTL1678 trans*-complemented with a 6×histidine-tagged FTL1678. Whole-cell lysates (WCL), OM fractions, and IM fractions were separated by SDS-PAGE, transferred to nitrocellulose, and immunoblotting was performed using antisera specific for the periplasmic protein TolB (αTolB).

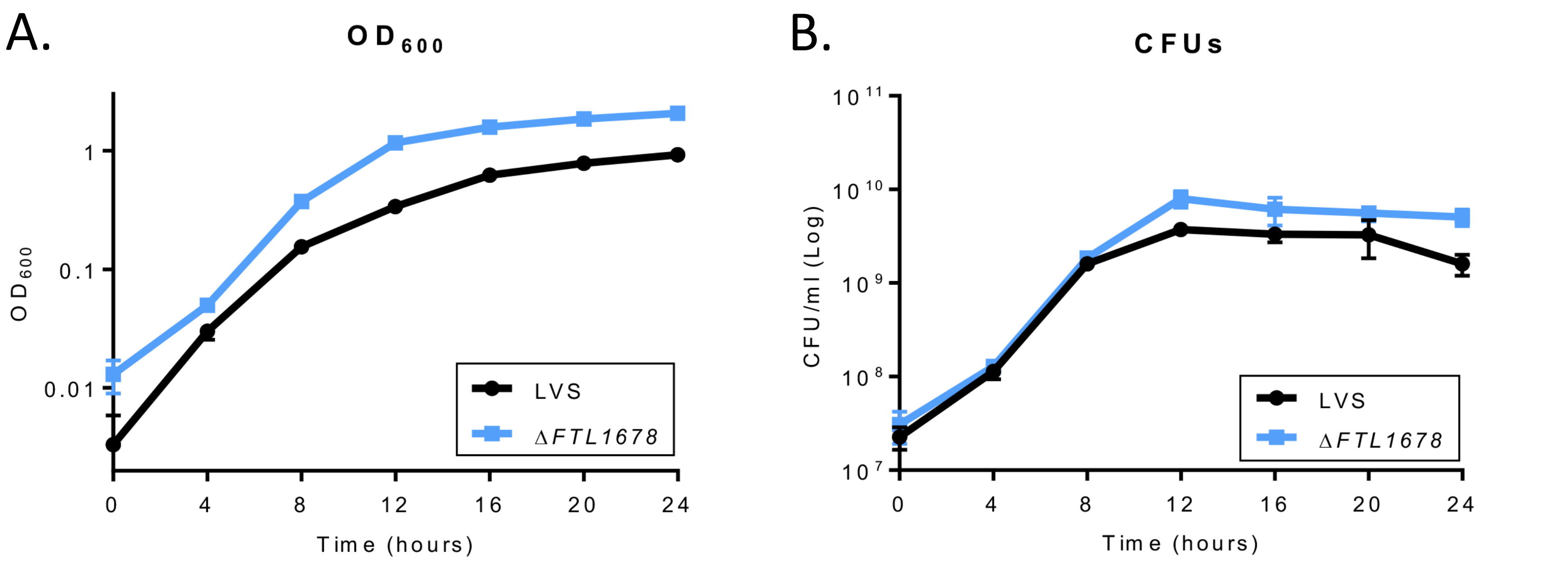


**Figure S5. Δ*FTL1678* does not have a growth defect.** WT and Δ*FTL1678* were grown in sMHB for 24 h at 37˚C. Samples were taken every 4 h for: (A) $OD_{600}$ measurements, and (B) CFU enumeration following serial-dilution and plating.

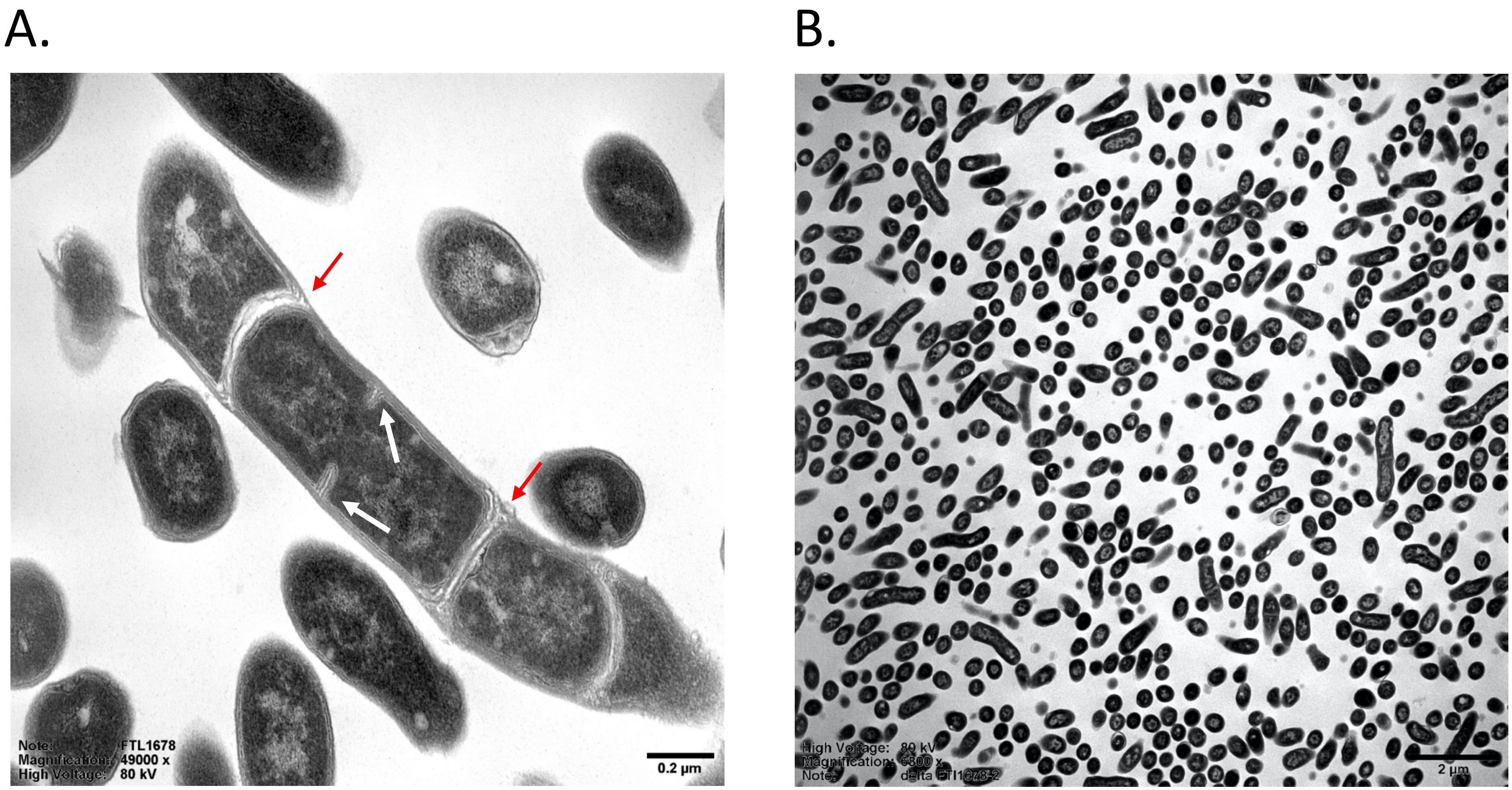


**Figure S6. Δ*FTL1678* has septation defects.** Transmission electron micrograph images of Δ*FTL1678* showing aberrant septal formation and reduced ability to separate cells. Images taken at: (A) 49,000×, scale bar represents 200 nm; and (B) 6,800×, scale bar represents 2 µm. In (A), red arrows point to formed septa that have not separated in Δ*FTL1678* and white arrows point to new septa that are forming in Δ*FTL1678*. Experiments were performed twice to confirm reproducibility, with two bacterial preparations fixed, stained, embedded, sectioned, and visualized per experiment. Representative images shown.

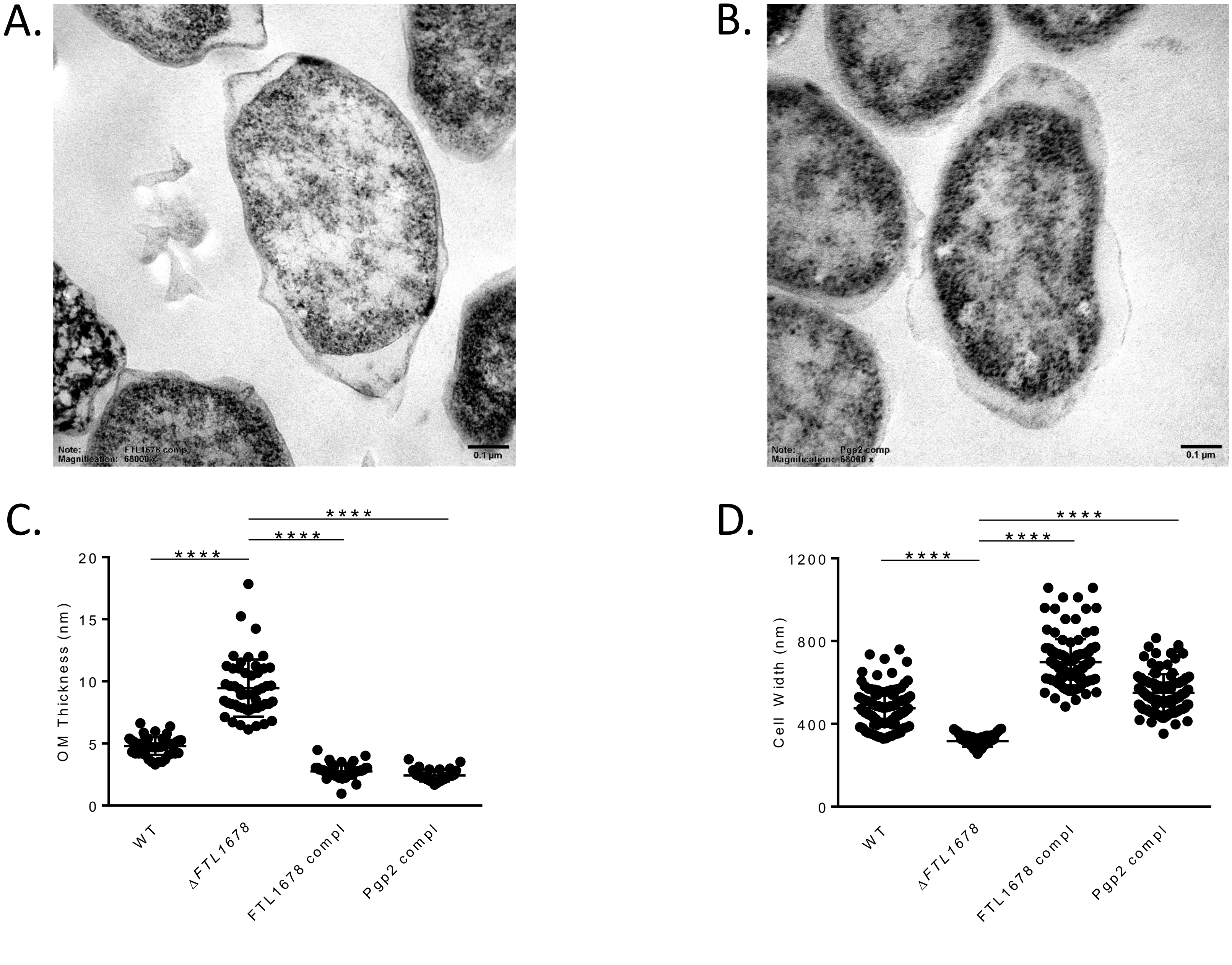
A.
Note: FTL1678 comp
Magnification: 68000 x
0.1 µm
B.
Note: Pgp2 comp
Magnification: 68000 x
0.1 µm
C.
OM Thickness (nm)
20
15
10
5
0
****
****
****
WT
ΔFTL1678
FTL1678 compl
Pgp2 compl
D.
Cell Width (nm)
1200
800
400
0
****
****
****
WT
ΔFTL1678
FTL1678 compl
Pgp2 compl

**Figure S7. FTL1678 *trans*-complement and *C. jejuni* Pgp2 *trans*-complement restore *F. tularensis* phenotype.** Transmission electron micrograph images of: (A) Δ*FTL1678 trans*-complemented with *FTL1678* [FTL1678 compl] or (B) Δ*FTL1678 trans*-complemented with *C. jejuni pgp2* [Pgp2 compl]. Bacteria were grown in sMHB to $OD_{600}$ of 0.4. Scale bars represent 100 nm. (C) Outer membrane thickness [WT n=50; Δ*FTL1678* n=50; FTL1678 compl n=44; Pgp2 compl n=33] and (D) cell width [WT n=119; Δ*FTL1678* n=120; FTL1678 compl n=119; Pgp2 compl n=119] of FTL1678 compl and Pgp2 compl were compared to WT LVS and Δ*FTL1678.* **** indicates $P < 0.0001$.

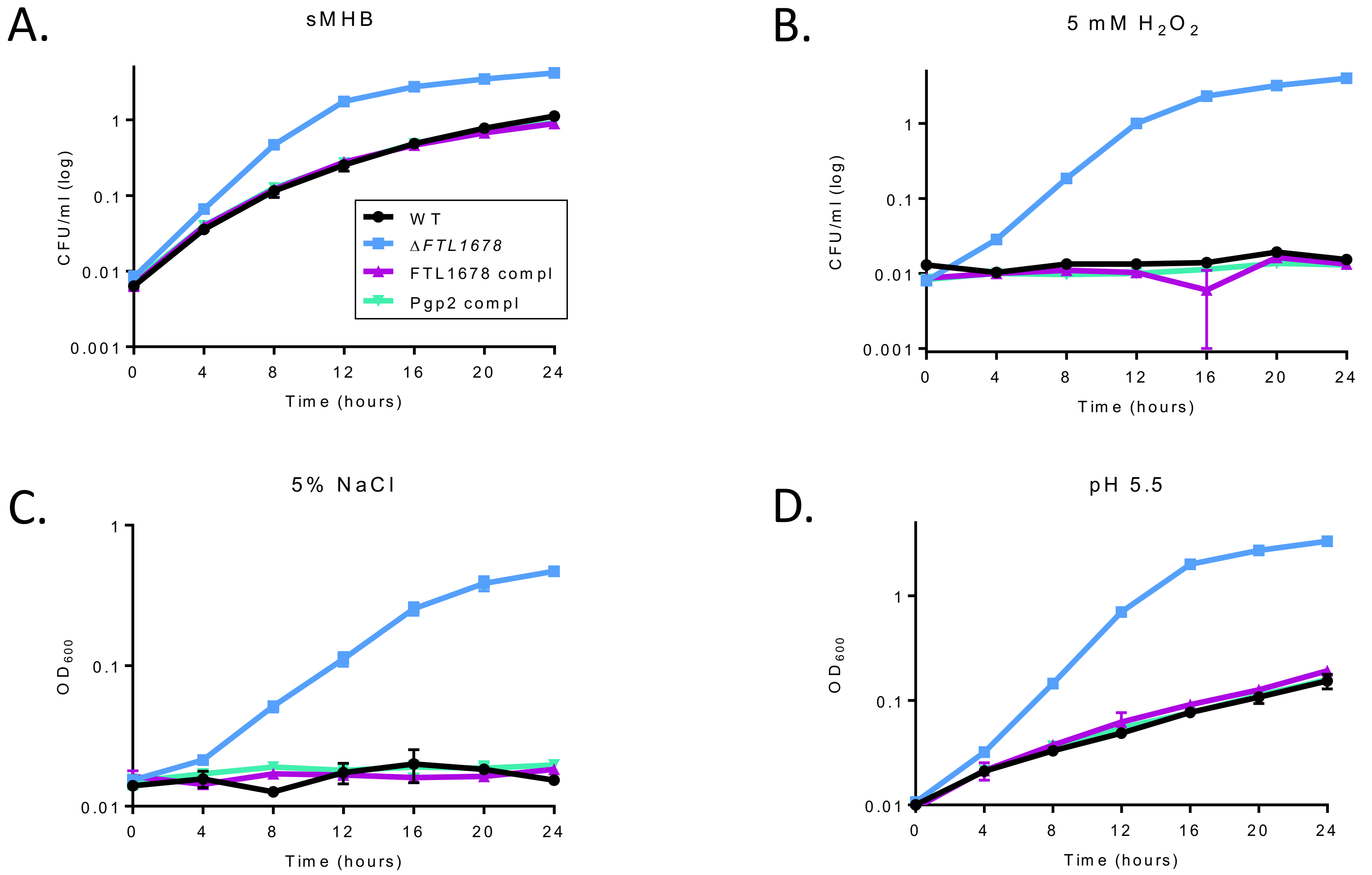
A.
sMHB
CFU/ml (log)
0.001
0.01
0.1
1
0
4
8
12
16
20
24
Time (hours)
WT
ΔFTL1678
FTL1678 compl
Pgp2 compl
B.
5 mM H2O2
CFU/ml (log)
0.001
0.01
0.1
1
0
4
8
12
16
20
24
Time (hours)
C.
5% NaCl
OD600
0.01
0.1
1
0
4
8
12
16
20
24
Time (hours)
D.
pH 5.5
OD600
0.01
0.1
1
0
4
8
12
16
20
24
Time (hours)

**Figure S8. FTL1678 *trans*-complement and *C. jejuni* Pgp2 *trans*-complement exhibit similar phenotypes to stressors as WT *F. tularensis*.** WT LVS [WT], Δ*FTL1678,* Δ*FTL1678 trans*-complemented with *FTL1678* [FTL1678 compl], or Δ*FTL1678 trans*-complemented with *C. jejuni pgp2* [Pgp2 compl] were grown in either: (A) sMHB; (B) sMHB with 5 mM $H_2O_2$; (C) sMHB with 5% NaCl; or (D) sMHB at pH 5.5. Cultures were incubated for 24 h and $OD_{600}$ measurements were recorded every 4 h.

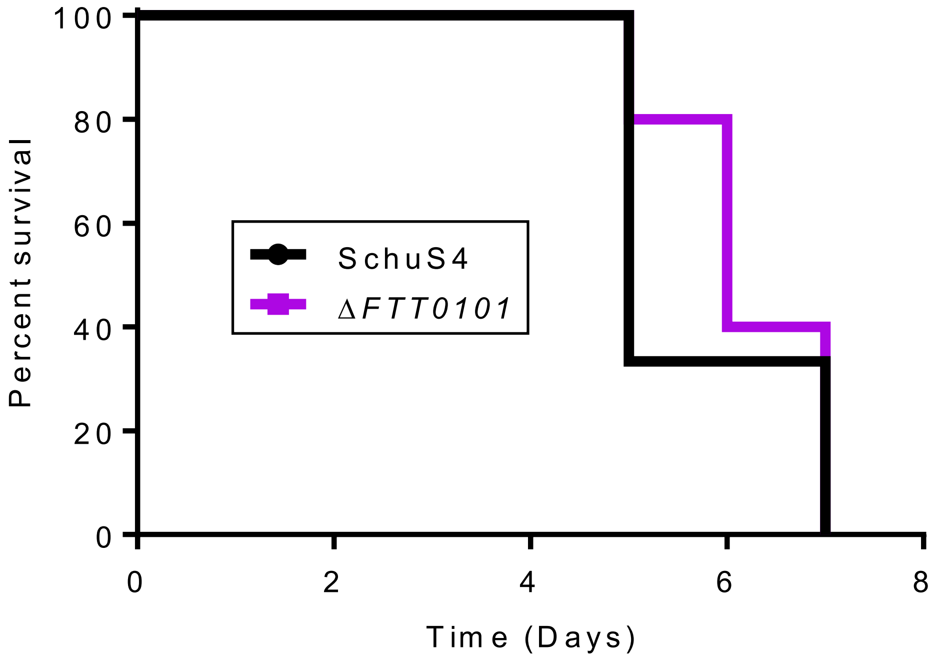


**Figure S9. FTT0101 is not required for *F. tularensis* Type A strain SchuS4 virulence.** C3H/HeN mice were intranasally infected with either 80 CFU SchuS4 (n=3 mice) or 12 CFU Δ*FTT0101* (n=5 mice).

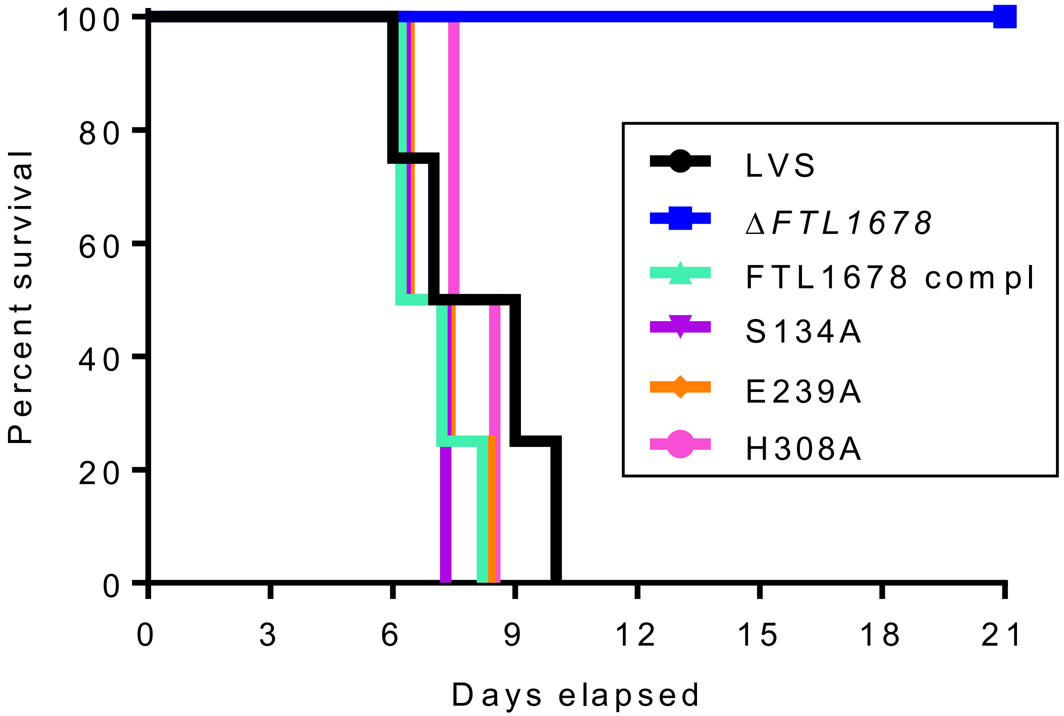


**Figure S10. Individual amino acids of the LdcA catalytic triad are not essential for *F. tularensis* virulence.** Groups of 5 C3H/HeN mice were intranasally-infected with $10^5$ CFU of either *F. tularensis* WT LVS, Δ*FTL1678*, Δ*FTL1678 trans*-complemented with FTL1678 (FTL1678 compl), Δ*FTL1678 trans*-complemented with S134A (S134A), Δ*FTL1678 trans*-complemented with E239A (E239A), or Δ*FTL1678 trans*-complemented with H308A (H308A). Animal health was monitored daily through day 21 post-infection. ** *P* < 0.01.
